# Insight into Lipopolysaccharide Translocation by Cryo-EM structures of a LptDE Transporter in Complex with Pro-Macrobodies

**DOI:** 10.1101/2021.03.23.436624

**Authors:** Mathieu Botte, Dongchun Ni, Stephan Schenck, Iwan Zimmermann, Mohamed Chami, Nicolas Bocquet, Pascal Egloff, Denis Bucher, Matilde Trabuco, Robert K.Y. Cheng, Janine D. Brunner, Markus A. Seeger, Henning Stahlberg, Michael Hennig

## Abstract

Lipopolysaccharides (LPS) are major constituents of the extracellular leaflet in the bacterial outer membrane and form an effective physical barrier for environmental threats and for antibiotics in Gram-negative bacteria ^1^. The last step of LPS insertion via the Lpt pathway is mediated by the LptD/E protein complex ^2^. Despite detailed insights from X-ray crystallography into the architecture of LptDE transporter complexes ^3–5^, no structure of a laterally open LptD transporter has been described, a transient state that occurs during LPS release ^6^. To facilitate the acquisition of hitherto unknown conformations we subjected LptDE of *N. gonorrhoeae* to cryo-EM analyses. In complex with newly designed rigid chaperones derived from nanobodies (Pro-Macrobodies, PMbs) we obtained a map of a partially opened LptDE transporter at 3.4 Å resolution and in addition we captured a laterally fully opened LptDE complex from a subset of particles. Our work offers new insights into the mechanism of LPS insertion, provides a structural framework for the development of antibiotics targeting LptD and describes a novel, highly rigid and widely applicable chaperone scaffold to enable structural biology of challenging protein targets.

## Introduction

Multi-drug resistant bacteria present a growing concern for human health ^7^. Among these, Gram-negative bacteria are well-shielded from their environment by an outer membrane (OM) that establishes a tight barrier for several antibiotics due to the high density of lipopolysaccharides (LPS) in the outer leaflet of the membrane. LPS are thus an essential component of the bacterial resistance. LPS is synthesized in the cytosol and transported across the periplasmic space to the outer leaflet via the Lpt pathway, consisting of the proteins LptA, B, C, D, E, F and G, which form a trans-envelope complex to bridge inner and outer membranes ^8–11^. Hence, disrupting the assembly of the outer membrane by inhibiting the Lpt pathway is an attractive strategy for novel antibiotic therapeutics ^12,13^. The membrane-integral LptDE complex in the OM executes the last step of LPS transport and is amongst other OM protein constituents a promising antibiotic target also due to its surface exposed localization^14,15^. The structures of LptDE from multiple species were determined by X-ray crystallography and provided detailed insight into the general architecture and the LPS path ^3–5^. The lateral opening of the LptD β-barrel, which enables the exit of LPS into the outer leaflet, has been conclusively inferred from simulations and mutagenesis studies ^16,17^, yet awaits structural evidence. From a pharmaceutical perspective, open conformations of barrel architectures are of high interest because of the propensity of β-hairpin mimetics (currently the most promising class of antibiotics to interfere with OM assembly proteins ^14,15,18^) to target to terminal β-strands by β-augmentation ^19^. Thus, more insight into the conformational space of LptD is in demand not only for an understanding of LPS transport in general but also for structure-based drug design. Here, we present the first cryo-EM structures of the LptDE transporter of the pathogen *Neisseria gonorrhoeae* (NgLptDE) with partially and fully opened lateral gates. Importantly, we could obtain these structures by complexation with nanobody-based chaperones. Departing from the structure of the previously described Macrobodies ^20^ that are built of a nanobody (Nb) and a C-terminally fused maltose-binding protein (MBP), we increased the rigidity of the original linker between the two moieties substantially to design a new chaperone scaffold, named Pro-Macrobodies, with excellent properties for improved particle classification and particle enlargement in cryo-EM.

## Results

### Design of Pro-Macrobodies and complexation with NgLptDE for cryo-EM analysis

To structurally characterize LptDE of *N. gonorrhoeae* and open the possibility of finding new conformations of these transporters we used single particle cryo electron microscopy (cryo-EM) as this method provides an opportunity to capture the conformational breadth of protein samples ^21–23^. After extensive optimization, we obtained a high-resolution structure of NgLptDE at 3.4 Å in complex with the enlarged synthetic VHHs PMb_21_ and PMb_51_ (Fig. 1a,b; Suppl. Figs. 1a-d, 4a-c). Our initial attempts to obtain a structure of uncomplexed NgLptDE by cryo-EM were limited to a resolution of 4.6 Å (Suppl. Fig. 1e-f). Consequently, molecular details such as side chains for confident tracing of the sequence for de novo building and the identification of potential ligands could not be visualized. During the course of our study we have generated sybodies (Sbs) which are synthetic nanobodies (Nbs) or VHHs ^24^ against NgLptDE (Suppl. Fig. 7a-c). We thought to transform the Sbs to macrobodies (Mbs) by fusion of MBP to the C-terminus as recently described for the crystallization of an ion channel ^20^ to use them as fiducial markers for improved particle classification. Since a chaperone for cryo-EM requires rigidity, the flexibility of Mb_51H01_ (extracted from the PDB entry 6HD8) (Fig. 2a) was initially assessed by molecular dynamics (MD) analysis. A bending motion of the MBP moiety was observed relative to the Nb of ~50° in multiple directions as well as torsional movements (Figs. 2a) due to high rotational freedom of the linker residues Val122 and Lys123 of Mb_51H01_ (Suppl. Fig. 2c). Despite their promising shape and size, Mbs are thus of limited use for particle enlargement in cryo-EM. *In silico* design of more rigid linkers was attempted. In particular, substitutions of Val122 and Lys123 by two consecutive prolines was found to lead to a stable chaperone and dampened motions as proline has the lowest rotational freedom of all amino acids. The mutated model was computationally predicted to adopt a new conformation, where the MBP moiety is turned by ~170 degrees according to the trans-configuration between Pro122 and Pro123 (Suppl. Fig. 2b). The mutated Mb, termed Pro-Macrobody (PMb), showed strongly reduced flexibility in MD simulations (Fig. 2a,d; Suppl. Fig. 2c) and retained the positive properties of Mbs such as their simple design and their elongated structure. We then transformed Sbs against NgLptDE into PMbs (Suppl. Fig. 2a) and crystallized PMb_21_. The X-ray structure of PMb_21_ at a resolution of 2 Å confirmed the predicted structure and showed the lowest B-factors around its linker region that forms a short poly-Pro helix II stretch (Fig. 2b,c; Suppl. Fig. 2d,e; Suppl. Table 2). The C-terminal end of the Sb-moiety and the N-terminal region of MBP feature rigid β-sheets that are only bridged by the intrinsically stiff di-proline linker without additional interactions that would stabilize interdomain motions (Suppl. Fig. 2e). The specific binding moiety could thus be exchanged without a change in the properties of the connection. Further, the PMbs retained their binding kinetics compared to the original Sbs using grating coupled interferometry (Suppl. Fig. 7c). Next, using only monomeric NgLptDE (Suppl. Fig. 3a,b; see below), we identified by size exclusion chromatography (SEC) a complex of NgLptDE with PMb_21_ and PMb_51_ (Suppl. Fig. 3c-e), two of the strongest binders from our analysis (Suppl. Fig. 7a,c). We subjected this quaternary complex to cryo-EM and obtained a map of high quality (Fig. 1b, Suppl. Table. 1, Suppl. Fig. 1c,d) significantly improved over the uncomplexed NgLptDE (Suppl. Fig. 1g,h). The rigidity of the PMbs is apparent from 2D-class averages (Fig. 2e) where the N- and C-terminal lobes of MBP are clearly recognizable. The respective identities of PMb_21_ and PMb_51_ could be assigned by 2D-class averages of a NgLptDE-PMb_21_ complex (Fig. 2e). PMb_21_ and PMb_51_ appear very similar in 2D-class averages in support of the universality and rigidity of the PMb-scaffold. When we subjected an NgLptDE complex with Mb_21_ and Mb_51_ (with the original Val-Lys linker) to cryo-EM it was apparent from 2D-class averages that these chaperones are more flexible and thus the MBP moiety is much less defined (Fig. 2e).

**Figure 1:**
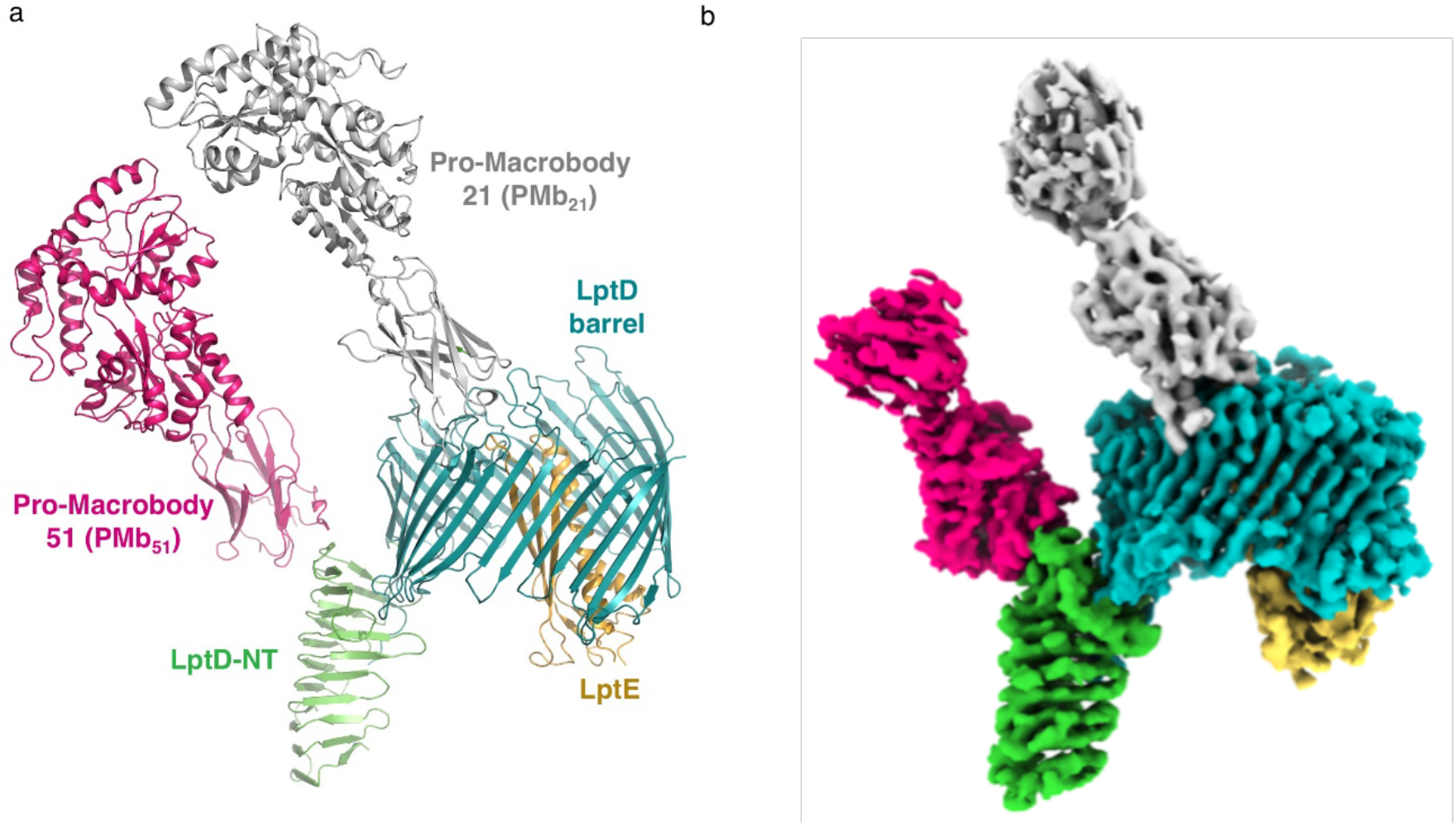
Cryo-EM structure of the NgLptDE-PMb_21_/PMb_51_ complex. (**a**) Side view of the NgLptDE-PMb_21_/PMb_51_ complex. (**b**) Cryo-EM map of the NgLptDE-PMb_21_/PMb_51_ complex at 3.4 Å. Colours encode domains as in (a).

**Figure 2:**
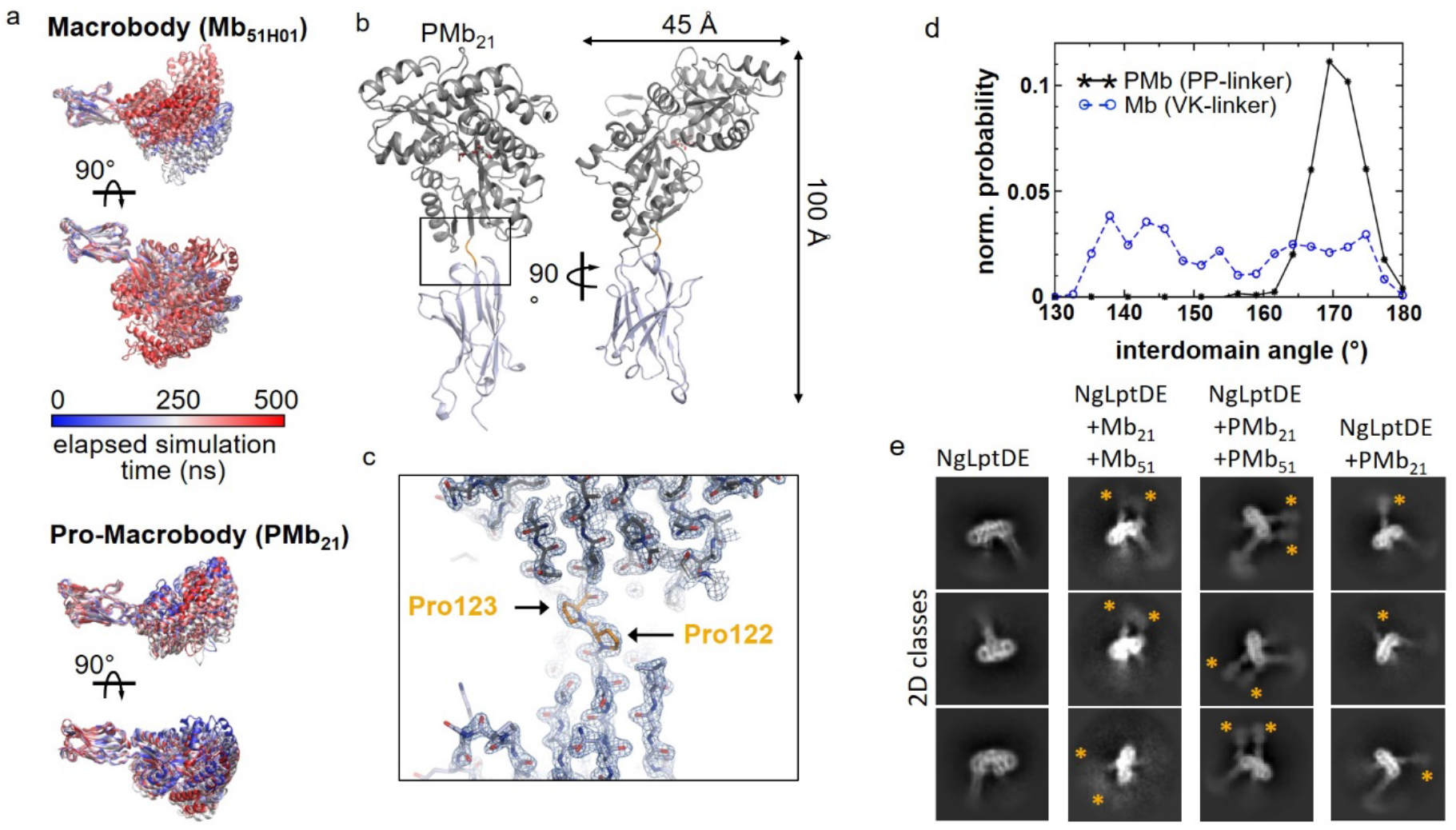
Generation and properties of Pro-Macrobodies. (**a**) Conformational flexibility between the Nb- and MBP-moieties connected by a Val-Lys linker (top) and Pro-Pro linker (bottom). Mb_51H01_ from PDB entry 6HD8 was subjected to a 500 ns all-atom MD simulation. Frames are colored by elapsed simulation time (middle panel) and aligned on the Nb-moiety to display the relative movements of MBP. The simulation was repeated with PMb_21_ using the X-ray structure shown in (**b**). (**c**) Enlarged view (boxed in (b) of the linker in the PMb_21_ crystal structure with the map contoured at (**d**) Plot of the normalized probability vs. the interdomain angle between the Nb and MBP in Mbs (Val-Lys linker) and PMbs (Pro-Pro linker) as derived from MD simulations. The PMbs show a defined distribution peaking at 170±5° whereas the original linker shows a wide distribution spanning almost 50°. (**e**) 2D-class averages of NgLptDE uncomplexed (left), complexed with Mbs 21 and 51 (left middle), complexed with PMbs 21 and 51 (middle right), and complexed with PMb_21_(right). The MBP moiety is indicated with yellow asterisks.

### The cryo-EM structure of NgLpTDE reveals a partially open LptD barrel

The cryo-EM structure of the complex showed the same overall LptDE architecture observed also in X-ray structures from LptDE of other Gram-negative bacteria ^25^ (Fig. 1a, Fig. 3a) but revealed also significant differences. Whereas NgLptE in our structure is nearly identical to LptE structures from other bacteria, our structure of NgLptD shows important deviations. The lateral and luminal gates in the cryo-EM structure of NgLptDE are more open compared to LptDE of *Klebsiella pneumoniae* (KpLptDE) and even more so compared to the X-ray structure of *Shigella flexneri* LptDE (SfLptDE) ^3–5^ (Fig. 4a-c). Only three hydrogen bonds were observed between β-sheets β1 and β26, which increases the separation of the two gating strands at their periplasmic end in NgLptDE by 3 Å compared to SfLptDE and KpLtDE, with at least six and five hydrogen bonds between those strands respectively (Figs. 4a-c). Luminal loop 1 preceding β1 adopts a conformation in NgLptDE that does not obstruct the luminal gate. Those residues of luminal loop 2 that are resolved in the cryo-EM map indicate an open conformation of this loop. This results in a direct connection between the hydrophobic groove of the N-terminal domain and the lumen of the β-barrel, which can be described as an opened luminal gate (Fig. 3a,b). The luminal gate has a diameter of approximately 10Å as determined by the smallest distance between Leu250-Asp251 and Asp768-Leu769, which, combined with an opened lateral gate, could allow passage of the LPS core and O-antigen from the periplasmic space to the lumen of the barrel. Overall, the NgLptDE structure shows a wider diameter of the barrel lumen reflecting a more open conformation of the luminal gate. With the distance between β1 and β26 being increased by 3 Å at the periplasmic side compare to SfLptD or KpLptD, this conformation represents a specific and until now unobserved stage of LPS transport.

**Figure 3:**
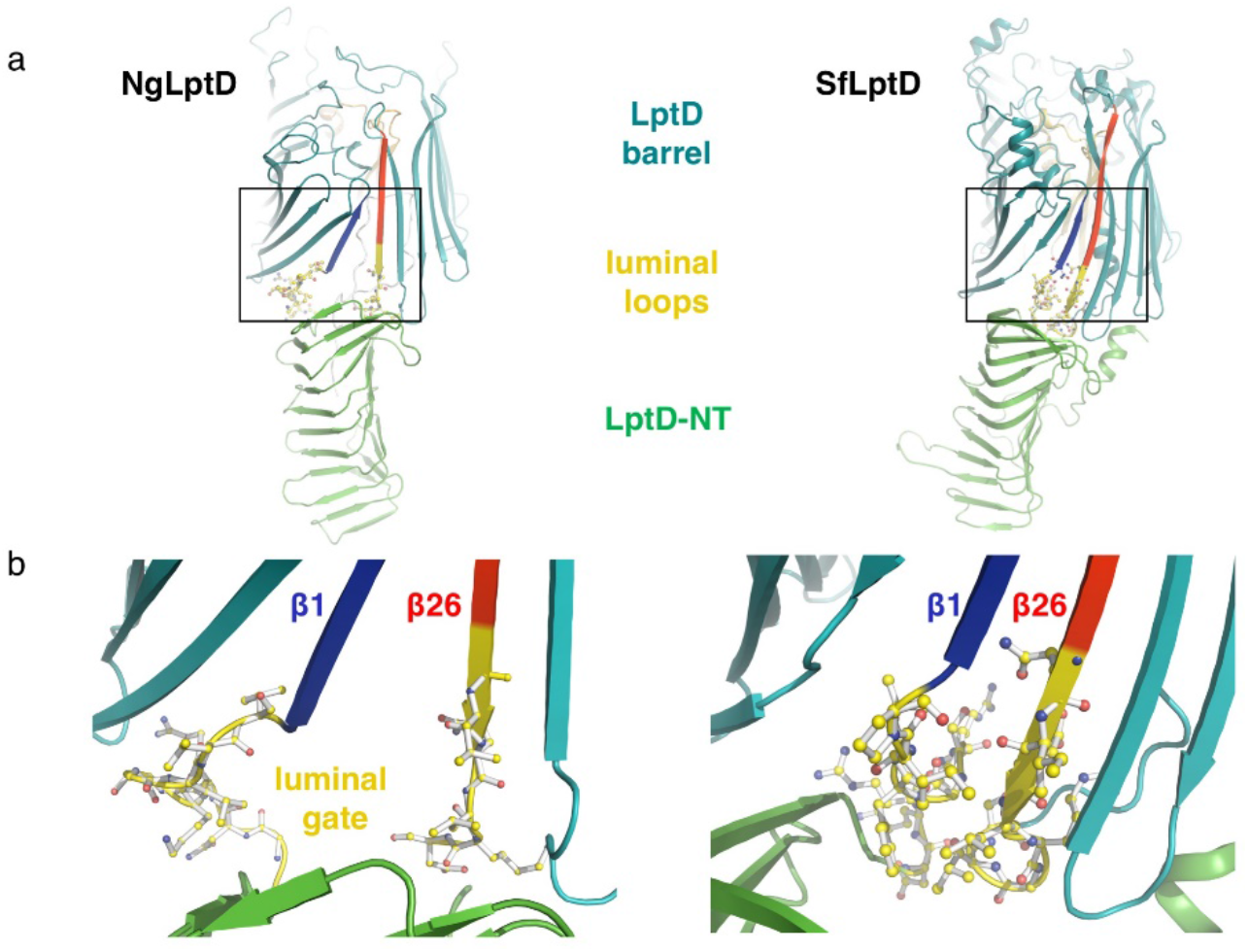
The luminal gate in structures of NgLptDE and SfLptDE. (**a**) The luminal loops (yellow) of NglptDE (left) are separated by more than 3 Å leading to a continuous groove between the LptD barrel and the hydrophobic groove of the N-terminal domain. In SfLptDE (PDB-ID 4Q35) (right) the luminal loops obstruct the luminal gate. (**b**) Enlarged view of the luminal gates of NgLptDE (left) and SfLptDE (right). The first (β1) and last (β26) strands of the LptD barrels are shown in blue and red respectively.

**Figure 4:**
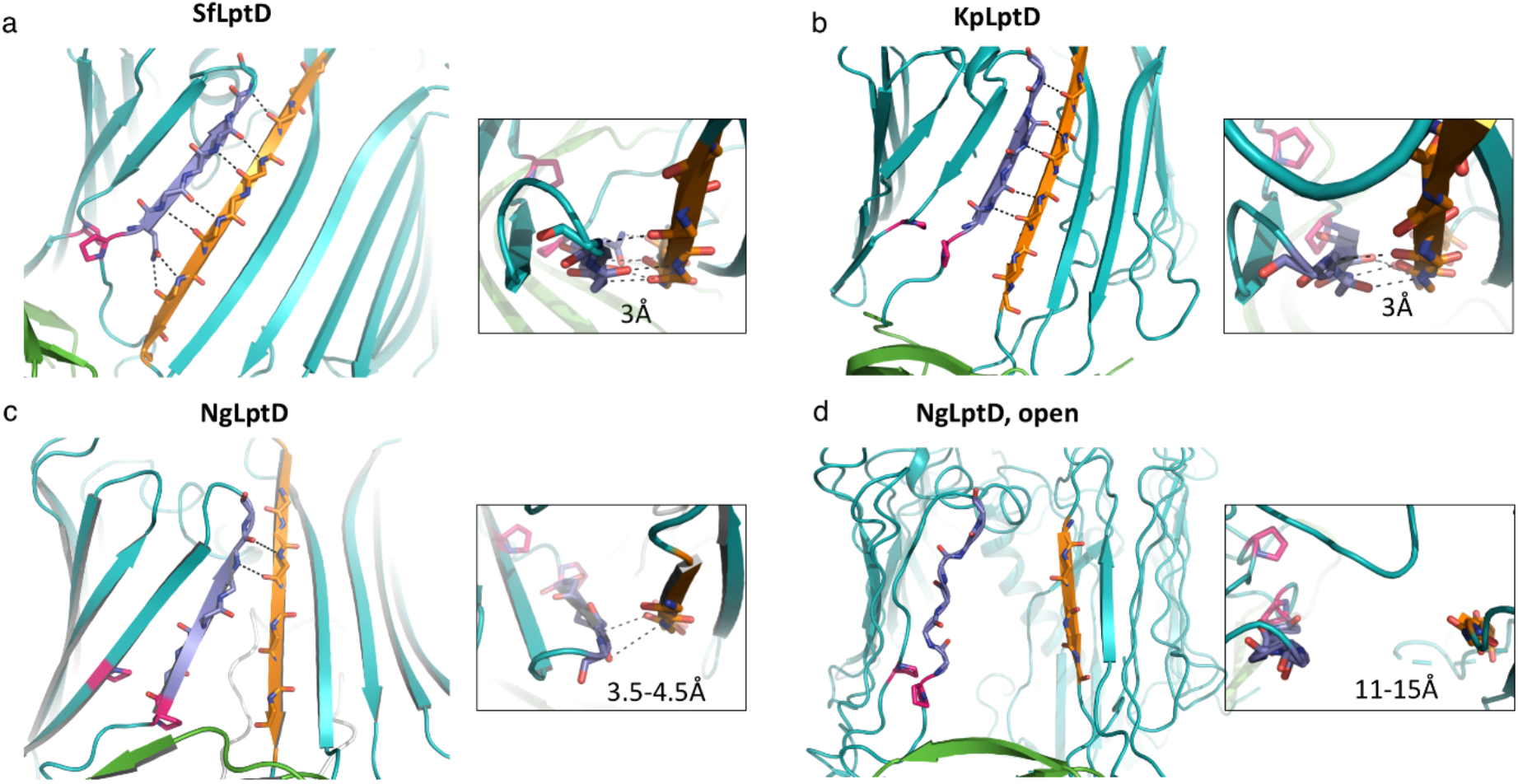
Hydrogen bonds between the β-strands 1 and 26 in structures of the LptD barrel. (**a**) SfLptDE, (**b**) KpLptDE (PDB-ID 5IV9), (**c**) NgLptDE (partially open), (**d**) NgLptDE (open lateral gate). Distances between the terminal strands are indicated and small insets show a view from extracellular. β-strand 1 and β26 are shown in light blue and orange respectively and the conserved proline residues of β-strands 1 and 2 in pink.

### Disulfide bridges in NgLptDE and PMb-binding sites

The N-terminus in our structure is not disulfide bonded to the periplasmic turn between β24 and β25 as in SfLptDE and KpLptDE, albeit a corresponding cysteine is present. The N-terminal 63 residues (not including the signal sequence) are therefore likely flexible and not visible in the cryo-EM map. The N-terminal domain is fixed by a disulfide bridge to the barrel, but differently from known full-length structures of KpLptDE and SfLptDE (Suppl. Fig. 5c). The conserved two consecutive cysteines at the turn between β24 and β25 are shifted by one position and separated by a glycine in contrast to most LptD proteins (Suppl. Fig. 5c). Folding of LptD proteins involves several steps of disulfide bond reorganization of which ultimately one disulfide bond is essential to connect the N-terminal jellyroll domain to the barrel ^26^. Despite these differences, the orientation of the N-terminal domain was the same as in KpLptDE, whereas that of SfLptdE is rotated by approximately 20° (Fig. 3a).

PMb_21_ binds partly to the barrel rim and to extracellular loops 5, 11, 12 and 13 that all have been described as dispensable for LptD function ^27^ (Suppl. Fig. 5a). PMb_51_ binds to the terminal β-strand of the jellyroll domain mainly via CDR3 that is forming a β-hairpin-like structure (β-strand augmentation) (Suppl. Fig. 5b). In the cell, this terminal strand of the jellyroll domain is deeply buried in the membrane such that PMb_51_ or its parent Sb_51_ could not interfere with LPS transport by partial blockade of the exit path. Both, Sb_21_ and Sb_51_ did not increase the susceptibility of *N. gonorrhoeae* towards vancomycin (Suppl. Fig. 7e). Further, neither of the Sbs bound to the surface of *E. coli* SF100 cells expressing NgLptDE (Suppl. Fig. 7f) which at least for Sb_51_ is not surprising due to the membrane localization of the epitope. Notably, paratopes like the CDR3 of PMb_51_ could also serve as template for the development of peptide antibiotics ^15^. It appears that the PMbs did not lock LptDE in a specific conformation as we observed additional states in the sample (see below).

### Additional density observed in NgLptD

The 25 C-terminal residues of the LptD barrel domain and luminal loop 2 were not visible in our initial cryo-EM map (“Overall” NgLptDE-PMb21-PMb51, Suppl. Table 1). In order to gain more insight into the function of the luminal gate in LPS transport, the cryo-EM data were re-analyzed focusing the refinement processing within a tight mask encompassing the N-terminal domain as well as β1-β26 region of NgLptD, only. The resulting cryo-EM map at 3.43 Å resolution allowed to trace the complete chain of the C-terminal region of LptD (Suppl. Fig. 6a-d). The C-terminal stretch following β26 extends deeply into the lumen of the β-barrel towards the restriction separating the two lobes of the barrel and in proximity to LptE. Interestingly, the C-terminal residues from Asn798 to Pro801 bind into the groove of the N-terminal jellyroll domain. Several salt bridges and hydrogen bonds stabilize the interaction (Suppl. Fig. 6d). Further work will be needed to determine if the observed position of the C-terminus could have a regulatory role similar to what has been suggested for the N-terminus ^4^. In contrast to the crystal structures of SfLptD and KpLptDE, no helical region is observed in luminal loop 2.

### A fully opened LptD barrel from a subset of particles

Image processing assigned about one fifth of the particles on the cryo-EM grid to a structure with an open barrel that did not fit the conformation of the main particle population. We used this subset of particles to compute a cryo-EM map of NgLptDE with an open lateral gate at 4.72 Å resolution using a very tight mask (Figs. 4d, 5a-c). This map shows a full opening of the β-barrel devoid of H-bonds between β1 and β26, and a separation of ~10Å between the extracellular ends and ~15Å between the periplasmic ends of these strands. The diameter of the open barrel was not significantly different to the one in the partly open conformation. Only the six N-terminal and four C-terminal strands were shifted by more than 1.5Å (Fig. 5b). This separation leads to a very large continuous solvent accessible channel from the extracellular space through the barrel to the periplasm that could easily accommodate transiting LPS molecules (Fig. 5a). Docking of LPS into this experimental structure indicated that the saccharide portion of LPS likely enters the barrel and passes through it to the extracellular face, while lipid A would be inserted into the outer membrane (Fig. 5c,d). These data provide structural evidence for previously suggested events that lead to strand separation and are compatible with the idea of hydrogen-bond weakening between strands β1 and β26 through conserved prolines in β1 and β2 ^5^. Further, the delineated path of the lipid from LPS-crosslinking experiments is in accord with the structure ^28^. Consequently, we propose that the structure of a laterally fully opened LptD barrel could serve as a geometrical guideline for the design of macrocyclic antibiotics.

**Figure 5:**
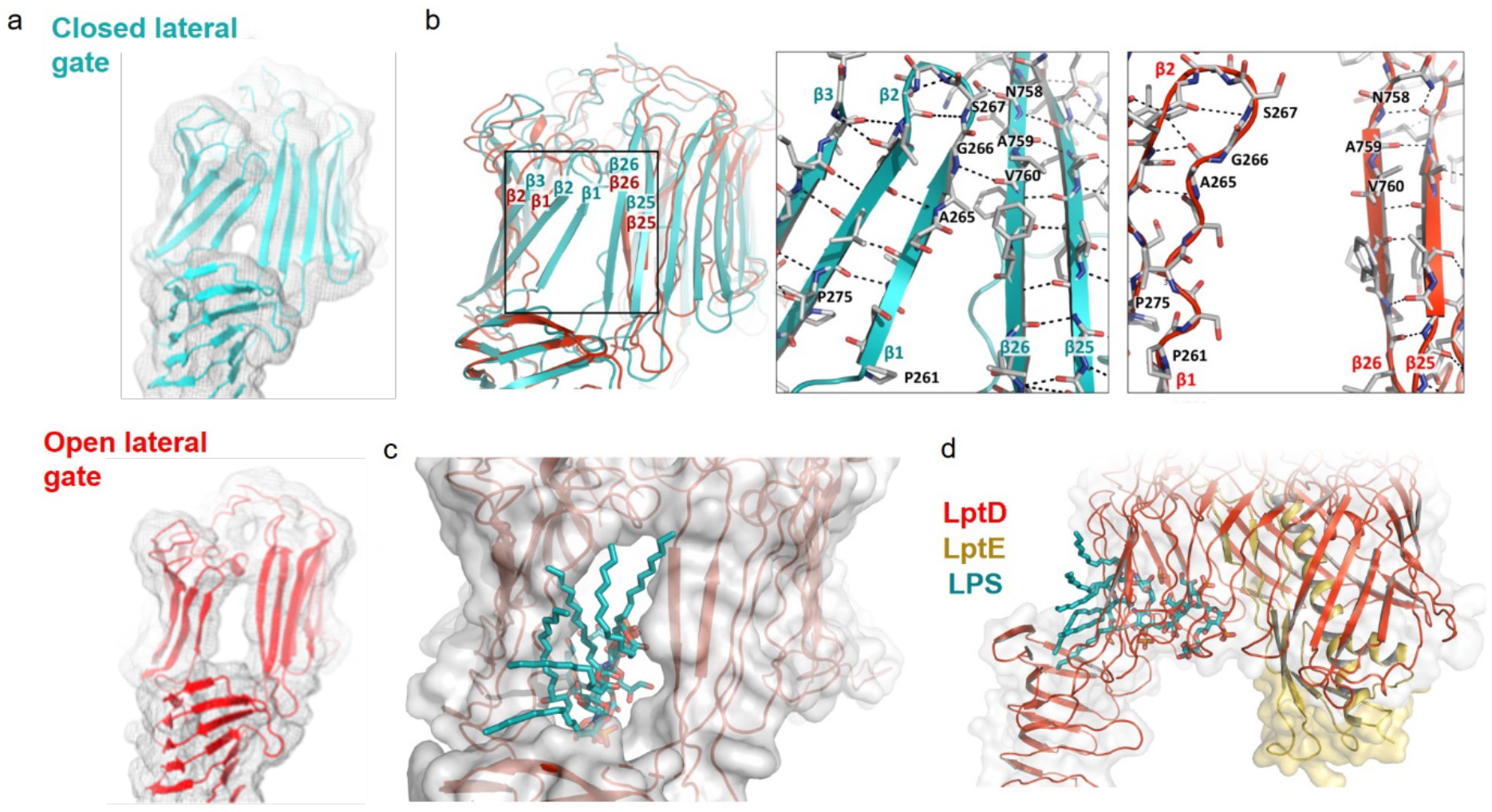
Comparison of the closed and open lateral gate of NgLptD. (**a**) Cryo-EM density of the NgLptD structures with closed (top) and open lateral gate (bottom) (maps displayed to the comparable contour level) with the respective model shown in cyan and red for the partly and fully open NgLptDE respectively (**b**) Superimposed models of NgLptD in partly (cyan) and fully open conformations (red). The boxes show the hydrogen-bond networks at the terminal strands of partly open (cyan, left) and fully open (red, right). (**c**) An LPS molecule (cyan) (extracted from 3FXI without modifications) fitted into the laterally open NgLptDE (red/yellow) viewed from extracellular or from the side (**d**).

### A dimer of NgLptDE mimics the assembly principle of the Lpt trans-envelope complex

The analysis of the cryo-EM data revealed a fraction of molecules is in a dimeric oligomeric state with a dimer interface right at the periplasmic end of the N-terminal LptD jelly-roll domain (Suppl. Fig. 8a-d). The dimerization was already observed in SEC analysis and turned out to be rather stable (Suppl. Fig. 3a,b). In the crystal lattices of KpLptDE and SfLptDE, molecular contacts via the N-terminal domain of LptD were observed as well. However, in our cryo-EM structure, the dimer was displaying a pseudo 2-fold symmetry and generated a continuous groove between the two beta jelly roll domains. This further corroborates the oligomerization of jelly-roll domains of the Lpt-pathway members LptD, LptA and LptC as a construction principle of the continuous hydrophobic slide across the periplasm ^5^. Presumably due to mobility in the N-terminal domain, applying a C2 symmetry for dimeric particles did not result in higher resolution (Suppl. Table 1).

## Discussion

### The structure of NgLptDE and implications for LPS transport

LPS transport requires a transient lateral opening of the LptD β-barrel to allow the synchronized translocation of the hydrophobic and hydrophilic portions of the large LPS molecules through the outer membrane^3–5^. For instance, cross-linking β-strands 1 and 26 through disulfide bridges abolished the function of LptD ^3,16^ providing clear indications that these strands separate at least transiently for LPS release. The strands 1 and 26 do furthermore show less hydrogen bonds compared to other barrel structures which support that these two strands are destined to temporarily separate. From mutagenesis studies and MD simulations it was proposed that the two conserved prolines Pro230 and Pro245 in *Yersinia pestis* LptD (corresponding to Pro261 and Pro275 in NgLptD, respectively) (Fig. 4) play a crucial role in opening ^5^ by attenuating the H-bonding between β1 and β26 to enable separation. Despite these clear indications, a lateral opening of the barrel was not fully evident from MD simulations ^5^ or if so, only in extreme regimes ^3^. Direct structural evidence for such a lateral opening was thus so far lacking. Using cryo-EM, we could find such a laterally opened structure that contributes to the completion of a full picture of LPS translocation. Our data suggests that the opening is a spontaneous event that does not strictly require a trigger, however simulations by computational methods suggest that the presence of LPS in the N-terminal domain promotes strand separation (i.e. increase the open-probability) ^17^. The precise regulation of barrel-opening thus requires further investigation, also with respect to the role of LptE ^29^ and the N-terminal domain of LptD ^30^.

### Implications for the development of new antibiotics

Closed β-strand architectures such as barrels offer naturally rather shallow binding sites, to modulate their function. The most promising antibiotics to target these proteins are currently β-hairpin mimetics (constrained macrocyclic peptides such as Pol 7080/ Murepavadin) that would bind to terminal beta-strands in open barrel intermediates or to components of the Lpt trans-envelope complex ^14,15,18,31^. Interestingly, thanatin, a naturally occurring antibiotically active peptide, targets to such β-strands of LptA, C and D ^30,32^. Non-hydrogen-bonded terminal β-strands which eventually appear temporarily at the seam of laterally open barrels or at the termini of LptA/C in the Lpt pathway can thus be understood as an Achilles` heel of these architectures due to their propensity to bind peptides or mimetics thereof. On LptD, known β-hairpin peptides target the N-terminus of the jelly-roll domain as shown recently ^30^. Taken together, the LptD structures of open barrel conformations could be particularly well-suited to serve as template for the design of new antibiotics. The cleft between β1 and β26 leaves sufficient space to accommodate a peptidomimetic that could bind through hydrogen-bonds to the LptD strands β1 and β26 (Suppl. Fig. 9). The cleft between β1 and β26 is an attractive ligand binding location since a peptide could bind on two strands, thereby increasing binding strength and specificity. The cleft’s localization at the extracellular space allows to expand the possibilities for peptide design to include also hydrophilic, charged and rather large residues as it does not require the passage through the lipidic portion of the OM.

A backfolding of the N-terminal loop to the N-terminal domain of LptD has been described in a previous study on SfLptDE ^4^. We observe for NgLptDE the C-terminus of LptD to be folded back to a very similar area on the N-terminal domain. It could thus be that LptD protein termini are involved in the regulation of LPS transport. Such regulatory areas could represent another potentially new site for a drug interaction inhibiting the LPS transport ^4,5^.

### Pro-Macrobodies as a chaperone for structural biology

Small membrane proteins still impose a significant challenge for structural analysis by cryo-EM and even more so β-barrel proteins because of the lower contrast provided in comparison to α-helical architectures. In these cases, binders that serve as fiducial markers can greatly improve resolution, especially for structure-based drug design to further advance. To date these chaperones are largely Fab fragments but PMbs described here, represent an attractive alternative for cryo-EM.

Pro-Macrobodies (PMbs) were an essential tool to elucidate the structure of the drug target NgLptDE by cryo-EM to high resolution. The complexation with the PMbs had positive impact on several parameters of the structural analysis by cryo-EM. A better randomization of particle orientations contributes to the higher resolution of complexed compared to uncomplexed NgLptDE (Suppl. Fig. 1b,f) as was described for the GABA_A_ receptor in complex with megabodies ^33,34^. For the EM analysis reported here, the major effect of PMbs is attributed to the increased size (230 kDa with PMbs vs. 110 kDa without) and the addition of distinct additional density to the particles for improved classification. Due to this enhanced particle selection and alignment, it was possible to reliably classify a sub-population of particles and find the laterally open conformation of NgLptD. PMbs are different from the recently described megabodies ^34^, as they feature only a single rigid linker. Our work provides evidence for a stable scaffold that significantly increased resolution of our target protein. Since any Nb or VHH can be converted to a PMb by fusing it in an identical rigid manner to the MBP moiety, they could become a widely applicable tool to improve results in cryo-EM, especially for small and challenging targets. This development encourages the application of cryo-EM to complement X-ray crystallography for structure-based drug discovery of novel medicines.

## Author contributions

M.B. and S.S. purified NgLptDE and NgLptDE-PMb complexes; M.B. and D.N. prepared samples for cryo-EM and collected data with M.C.; M.B., D.N. and H.S. analyzed cryo-EM data and built the models.; M.H. and M.A.S. wrote the grant for sybody generation. I.Z., P.E. and M.A.S. generated sybodies and performed and analyzed microbiological assays.; N.B. analyzed binding kinetics of NgLptDE-PMb and sybody complexes.; S.S., M.T. and R.K.Y.C. crystallized PMb21. M.T. and R.K.Y.C. solved the structure of PMb21 and built the model.; S.S., J.D.B. and D.B. designed PMbs, performed MD simulations and purified PMbs.; M.B., S.S., M.A.S., J.D.B., H.S. and M.H. wrote the manuscript with input from all other authors.

## Acknowledgements

This work was supported by the Innosuisse grant 25864.1 PFLS-LS to leadXpro AG and M.A.S.. M.H. and M.B. would like to thank Professor Yihua Huang (National Laboratory of Biomacromolecules, National Center of Protein Science-Beijing, Institute of Biophysics, Chinese Academy of Sciences, Beijing 100101, China) for kindly providing expression bacterial strain and expression vector but most importantly for scientific discussion.

## Data availability

EM raw image data were deposited in the Electron Microscopy Public Image Archive (EMPIAR) under accession number EMPIAR-XXXX. EM maps were deposited in the Electron Microscopy Data Bank under accession codes EMD-XXXX (), EMD-XXXX(), EMD-XXXX(), EMD-XXXX() and EMD-XXXX(). Atomic coordinates for NgLptDE from the cryo-EM study were deposited in the Protein Data Bank under accession codes PDB-XXXX. The X-ray structure of PMb_21_ was deposited with accession codes PDB-XXXX. All other data are available from the corresponding authors upon reasonable request.

---

**Suppl. Figure 1:**
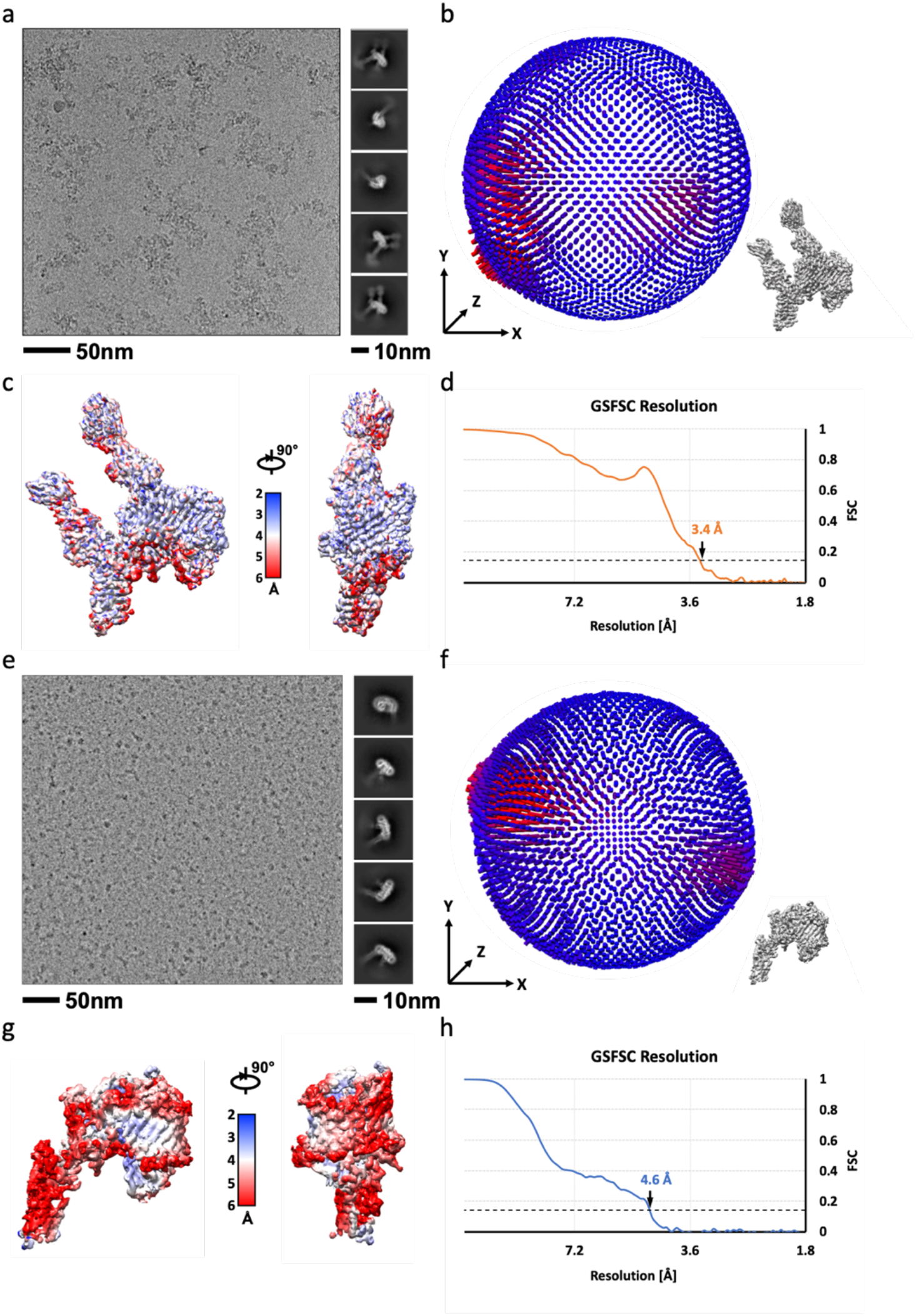
Cryo-EM acquisition and maps of complexed and uncomplexed NgLptDE. (**a**) Example micrograph of vitrified NgLptDE-PMb_21_/PMb_51_ quaternary complex. Class averages are shown on the right. (**b**) Distribution of the particle views that were used to build the map projected on a sphere relative to the orientation shown on the lower right. (**c**) Map of the laterally closed NgLptDE-PMb_21_/PMb_51_ quaternary complex with color-coded local resolution (middle panel). (**d**) Gold standard Fourier-Shell correlation resolution plot with a cutoff at 0.143 indicated by a dashed line. (**e-h**) Respective data as in (a-d) for the uncomplexed NgLptDE dataset.

**Suppl. Figure 2:**
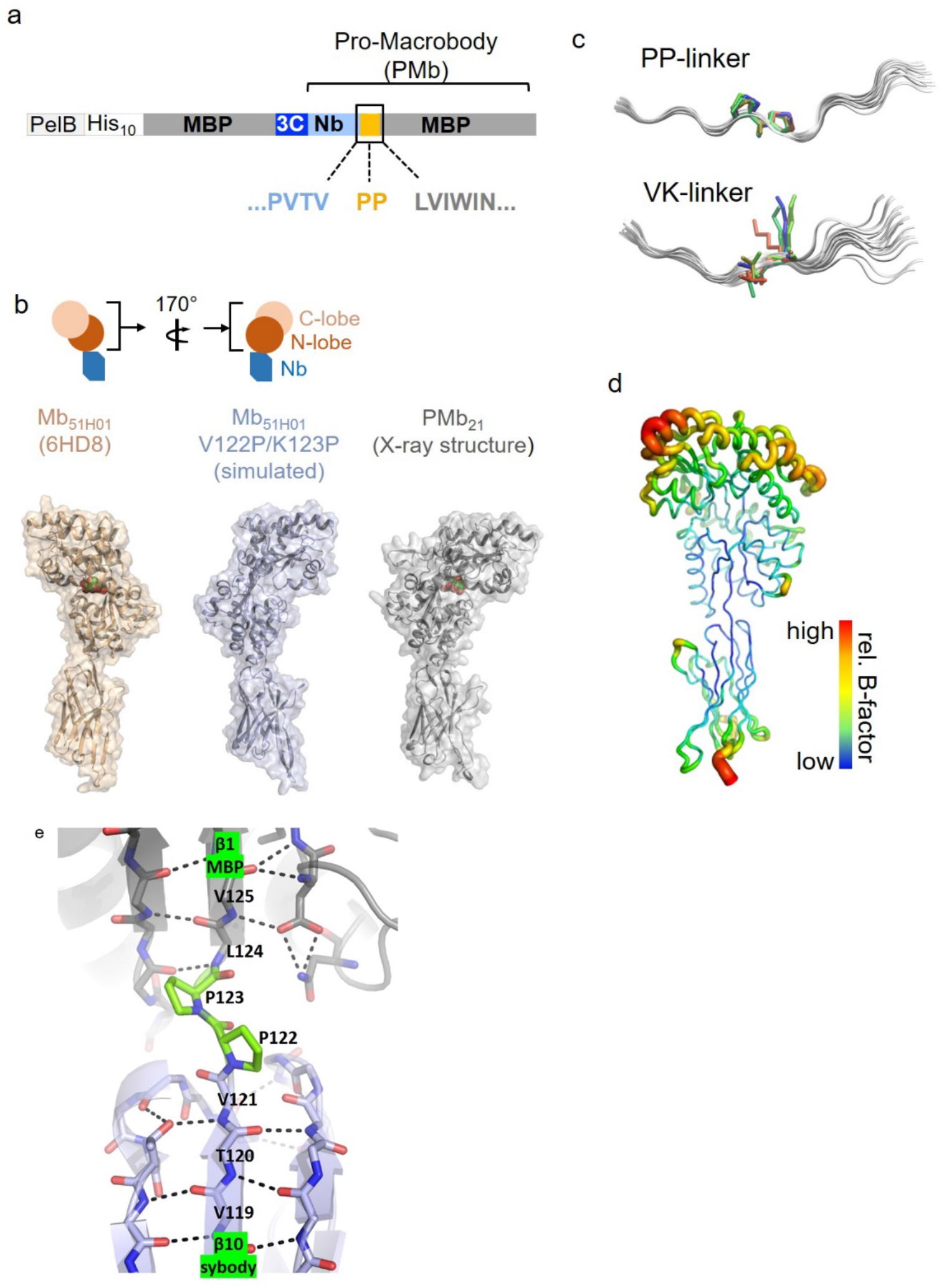
Design and properties of Pro-Macrobodies. (**a**) Construct layout for the expression of PMbs. (**b**) Comparison of the structures of the template Mb_51H01_ (left), a frame from the simulated (in-silico mutated) Mb_51H01_(middle) and the PMb_21_ X-ray structure. The orientation of the MBP moiety in Mb_51H01_ and its mutated version relative to the Nb-moiety is shown on top. Maltose is shown as space-fill model in the left and right panels. (**c**) C-alpha traces of the linker region in the MD simulations for the two types of linker. Single frames of the MD simulations were stacked, and the linker residues are shown as sticks. (**d**) Relative B-factors of the PMb_21_ crystal structure shown as putty in rainbow colors. (**e**) Hydrogen bonds around the linker region of PMb_21_ indicated by black dashes. The connected β-strands of the sybody (β10) and MBP (β1) are indicated together with the linker residues. The newly introduced residues P122/123 are labelled green. The C-terminal end of the sybody and the N-terminal part of MBP are dominated by β-sheets that are interrupted by the intrinsically rigid di-proline motif.

**Suppl. Figure 3:**
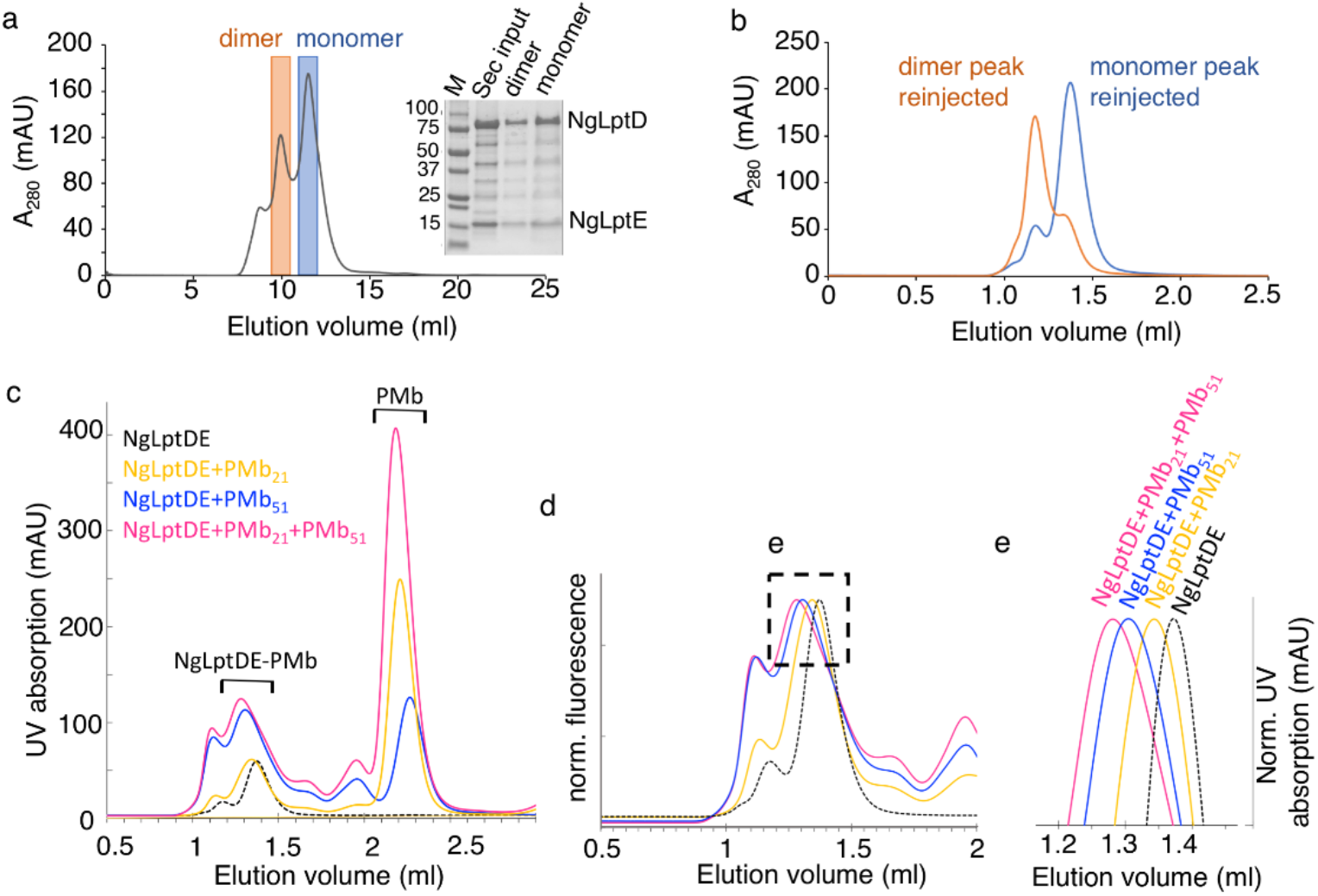
Size exclusion chromatography of NgLptDE and NgLptDE-PMb complexes. (**a**) Elution profile of dialysed and concentrated NgLptDE after IMAC subjected to a Superdex 200 10/300 Increase column. Dimer and Monomer peaks are indicated in orange and blue. A Coomassie stained SDS gel is shown as inset with LptD and LptE indicated. (**b**) Reinjection of the peak fractions with dimeric and monomeric NgLptDE to a Superdex 200 5/150 column. (**c**) NgLptDE mixed with excess of PMbs and subjected to a Superdex 200 5/150 column. Elution was monitored by Trp-fluorescence. (**d,e**) Enlarged and normalized peaks of NgLptDE and its complexes with PMbs in (**c**).

**Suppl. Figure 4:**
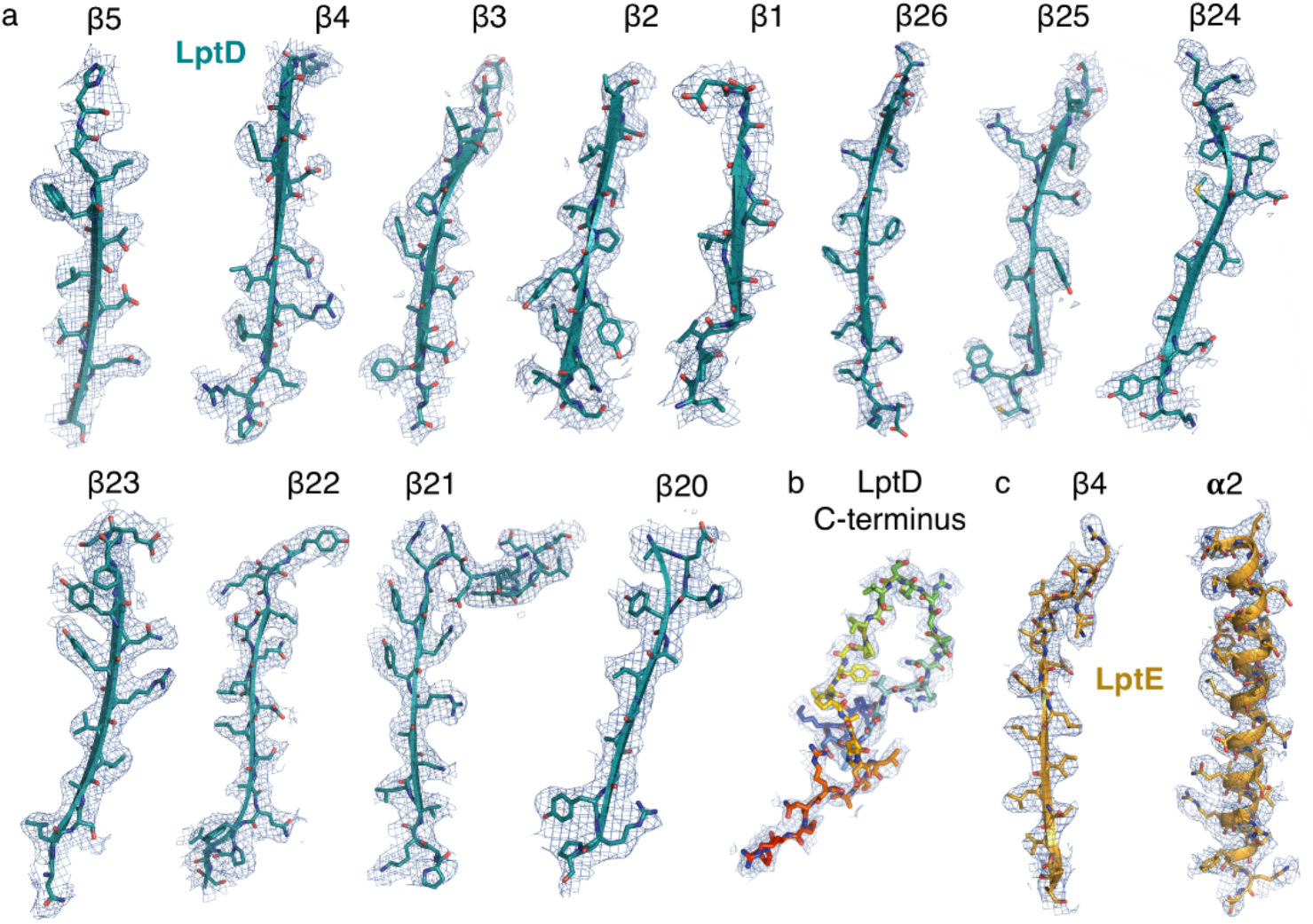
Agreement of the Cryo-EM map and the corresponding model. (**a**) β-strands of the NgLptD barrel shown with the cryo-EM map contoured at 9σ with the model superimposed. (**b**) LptD C-terminus colored in rainbow from N-to C-terminal ends. (**c**) β-strand 4 and α-helix 2 of LptE.

**Suppl. Figure 5:**
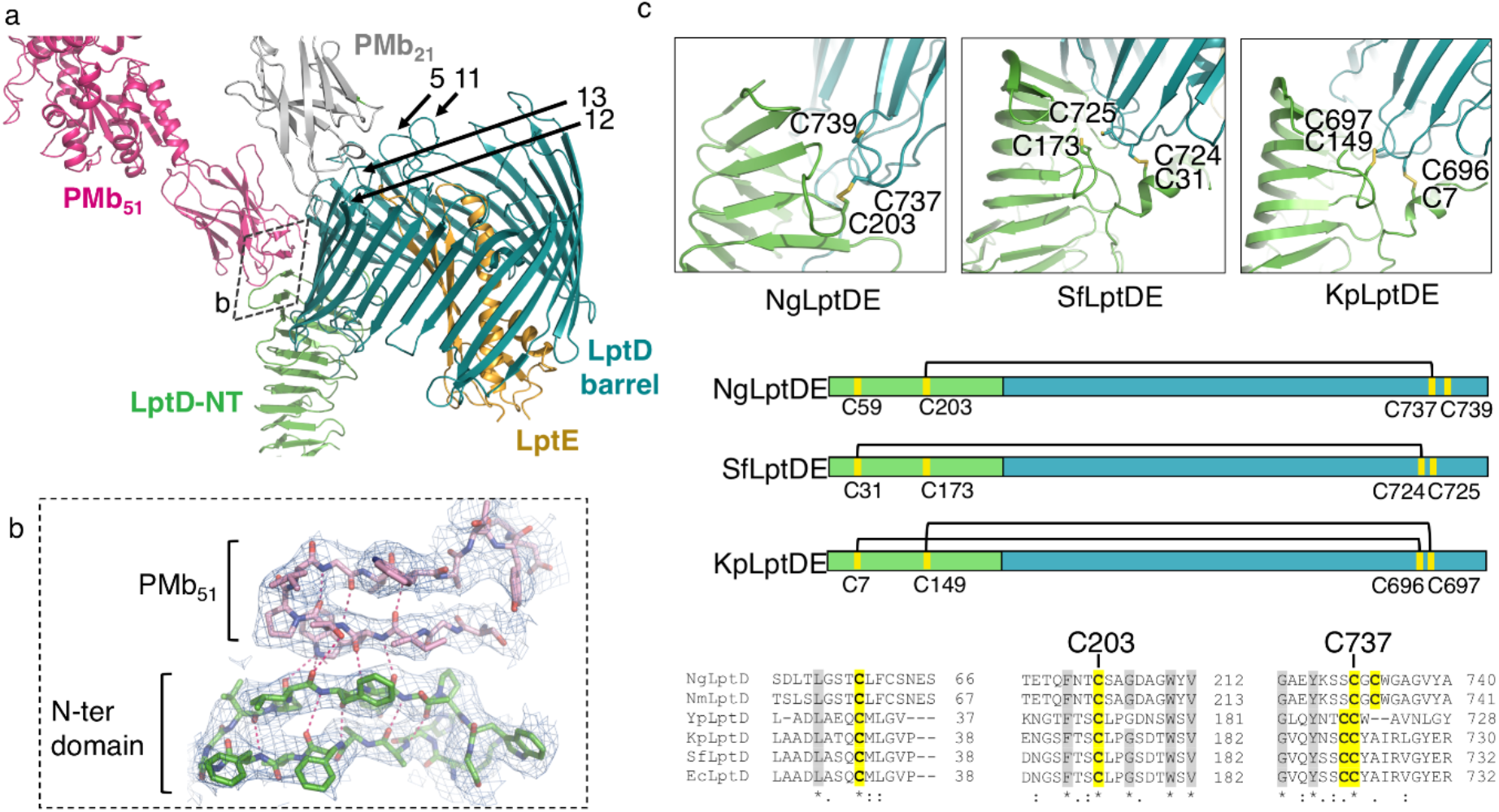
Details of the PMb51 CDR3 binding and disulfide bridges in LptD. (**a**) Overview of the binding regions of PMb_21_ and PMb_51_ at LptD barrel-loops and N-terminal domain. (**b**) View from the lateral seam in the LptD barrel (boxed in (a)) onto the edge of the N-terminal domain showing the hydrogen bonds between CDR3 and the terminal strand of the N-terminal domain and intramolecular bonds of CDR3. (**c**) Disulfide bridge patterns of conserved cysteines in LptD proteins from different bacterial species. The scheme in the middle shows the disulfide bridges schematically. An alignment (bottom) shows that the cysteines at the C-terminal end of *Neisseria* species are spaced by a glycine and thus different from other gram-negative bacteria.

**Suppl. Figure 6:**
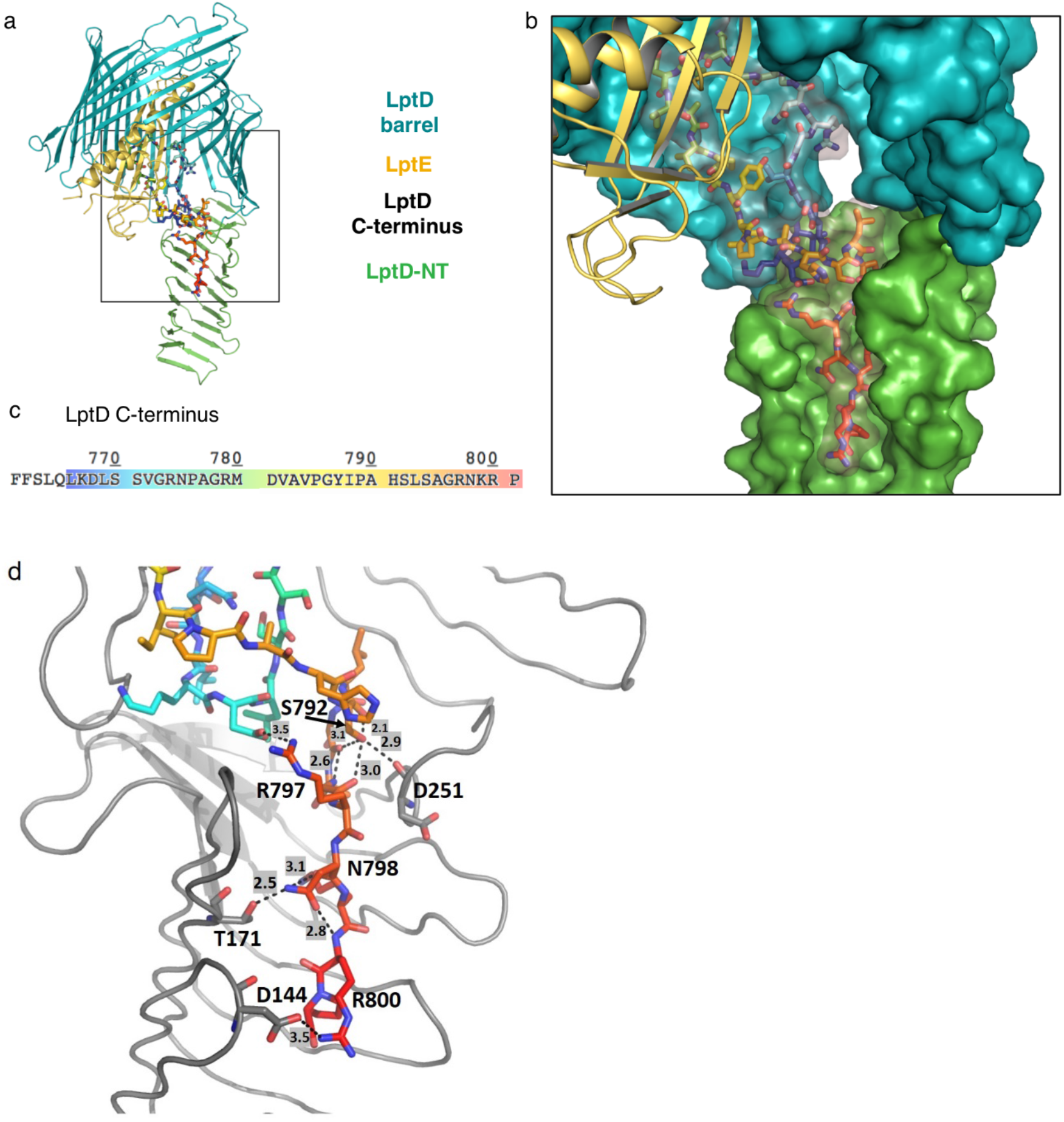
Binding of the C-terminus of NgLptD to the N-terminal jellyroll domain. (**a**) Overview of the location of the C-terminus in NgLptD in the complex. C-terminal residues are shown as sticks in rainbow colors as depicted in (c). (**b**) Close-up view of the C-terminus in the N-terminal domain in space fill model (transparent) with residues shown as sticks in rainbow colors. (**c**) Color code for the C-terminus used in (a,b,d) (**d**) Hydrogen bonds and salt bridges of the C-terminus (rainbow) with residues of the N-terminal domain (grey). Distances and involved residues are indicated.

**Suppl. Figure 7:**
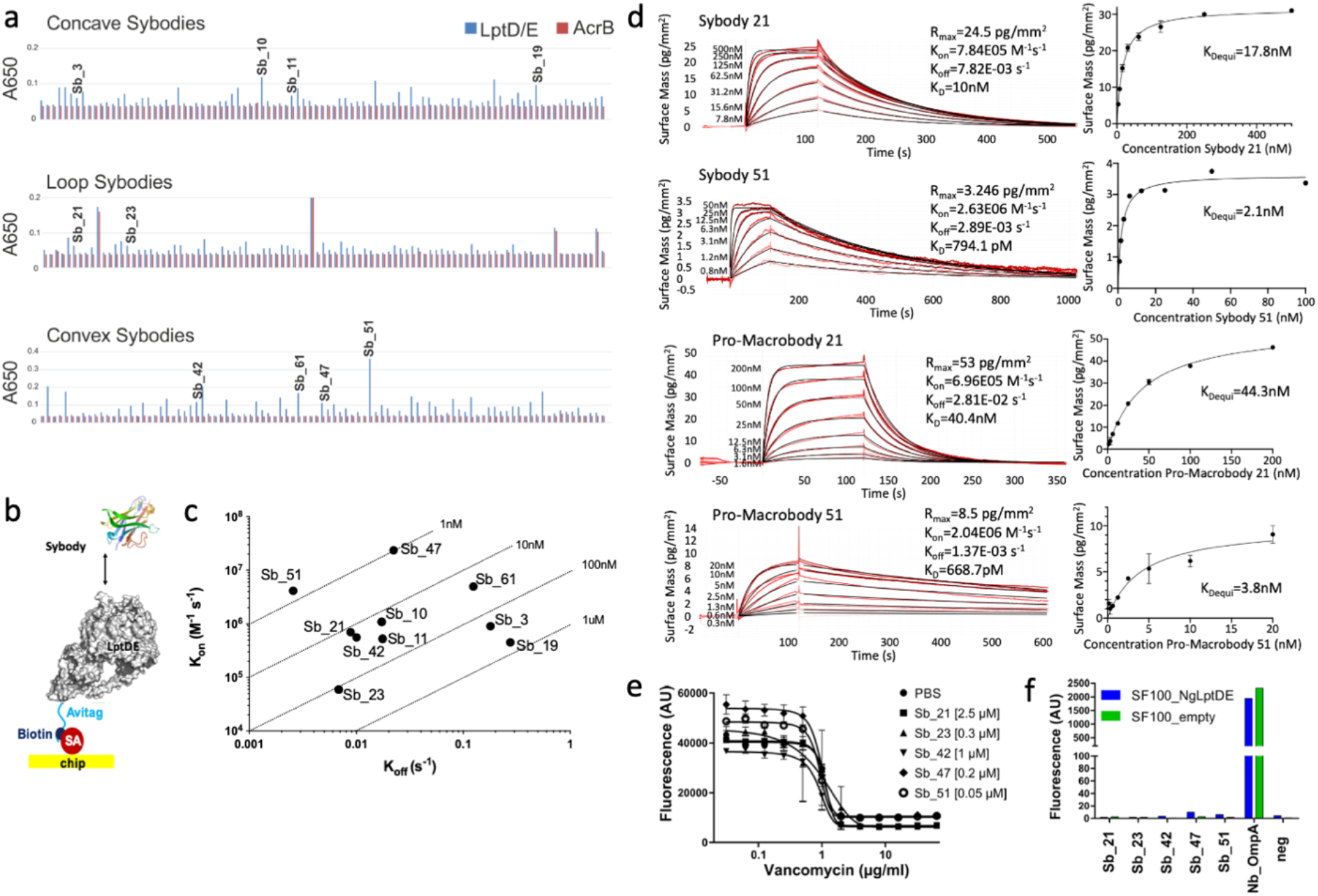
Sybody generation, characterization and comparison to PMb versions. (**a**) ELISA analysis of sybodies from the concave (top panel), loop (middle panel) and convex (lower panel) library. ELISA signals towards NgLptDE are shown in blue and control ELISA signals against AcrB are shown in red. Sybodies that were sequenced, purified and analysed by waveguide interferometry are indicated by providing the respective Sb_# numbers above the bars. (**b**) Illustration of the selection procedure using immobilized NgLptDE on Streptavidin chips as used for waveguide interferometry. (**c**) Evaluation of the binding kinetics of NgLptDE-positive sybodies using waveguide interferometry. (**d**) Binding kinetics of Sb21 and Sb51 (top panels) and PMb21/51 lower panels. KD values for Sb21, Sb51, PMb21 and PMb51 are 10 nM, 794.1 pM, 40.4 nM and 668.7 pM respectively. (**e**) Inhibition assay of *N. gonorrhoeae* growth in the presence of sybodies at concentrations indicated in the legend and varying concentrations of vancomycin. (**f**) Binding of fluorescently labeled sybodies on the surface of *E. coli* SF100 cells expressing NgLptDE (blue bars) or the same cells containing an empty vector control (green bars). Anti-OmpA Nbs are used as positive control (Nb_OmpA). No binders were added to the negative control (neg).

**Suppl. Figure 8:**
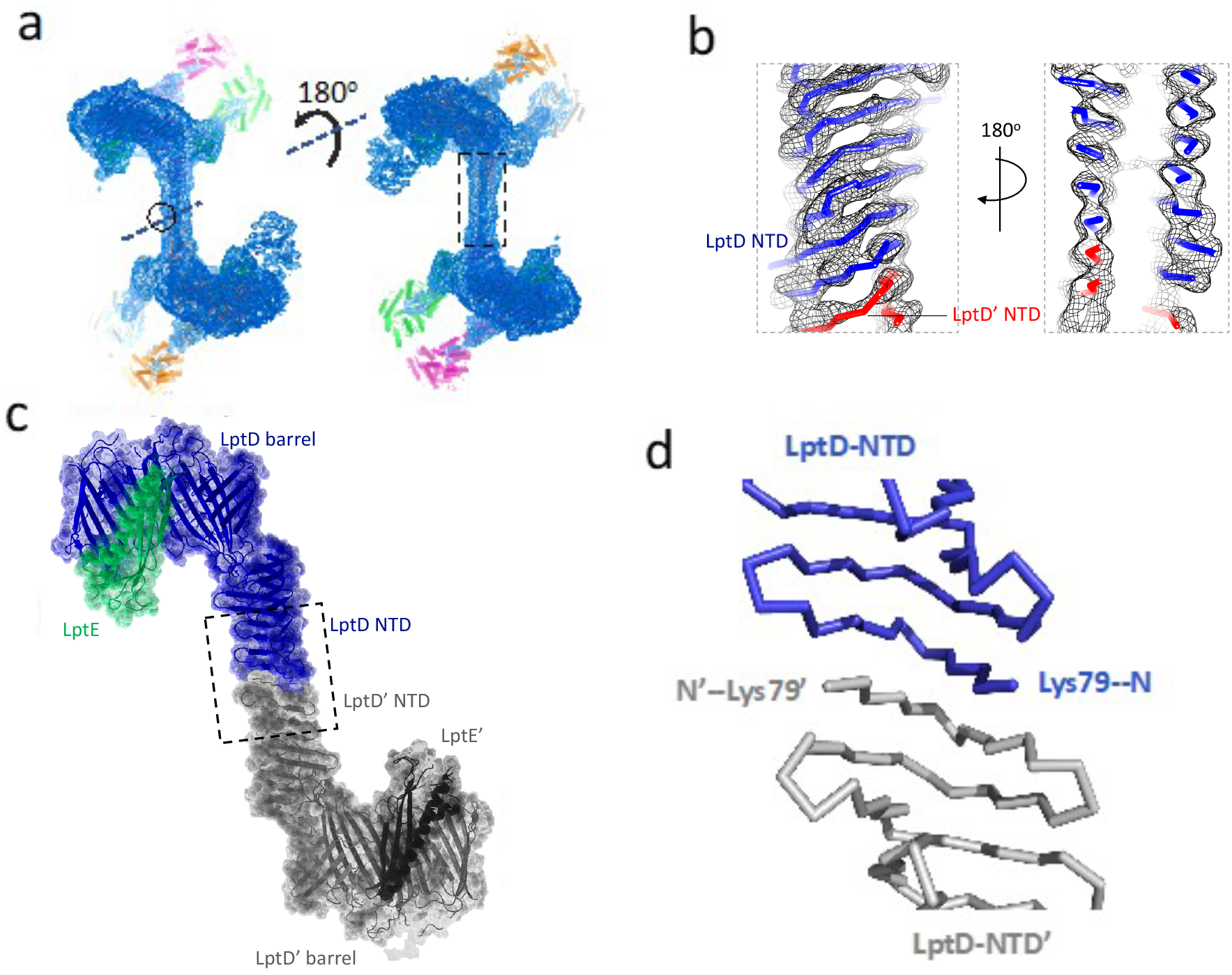
Head-to-head dimers of NgLptDE-PMb_21/51_ complexes. (**a**) Map of the head-to-head dimer of the NgLptDE-PMb_21/51_ complex. (**b**) Enlarged view of the stalk formed by the N-terminal regions boxed in (a). (**c**) Model of the dimer with enlarged view of the head-to-head dimerization by β-strand complementation shown in (**d**).

**Suppl. Figure 9:**
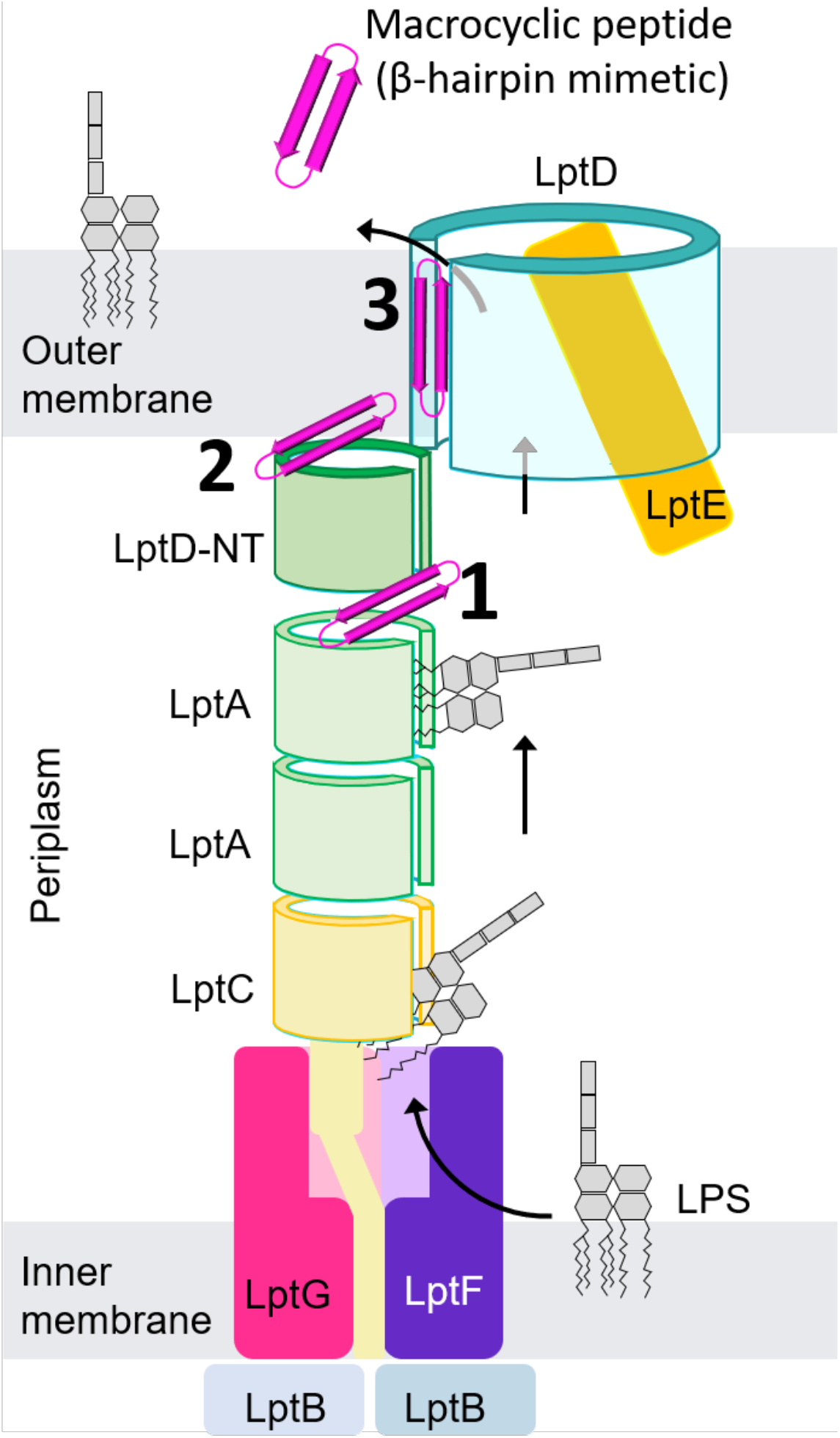
Binding sites for peptidomimetics targeting LptD. LPS is transported through the trans-envelope complex via the Lpt-pathway (black arrows). Blocking of this route by peptidomimetics that bind to terminal β-strands is a promising strategy to interfere with OM biogenesis of gram-negative pathogenic bacteria. Thanatin, a naturally occurring peptide, binds to the N-terminus of LptD next to described LptA interactions and disrupts the assembly of the LPS-route (1). Attractive binding sites for peptidomimetics are also present at the end of the N-terminal domain of LptD (2) like the β-augmentation by the CDR3 loop of PMb51 we observed in the structure and in particular in the cleft between β1 and β26 of a transiently open LptD barrel (3). The latter would be most accessible for targeting compounds as the OM would not need to be traversed.

**Suppl. table 1:** Cryo-EM data collection, refinement and statistics.

|  | NgLptDE-PMb<sub>21</sub>-PMb<sub>51</sub> |  |  |  |  | Apo NgLptDE |  |
| --- | --- | --- | --- | --- | --- | --- | --- |
|  |  |  |  |  |  | Monomeric fraction | Dimeric fraction |
| Microscope | FEI Titan Krios |  |  |  |  | FEI Titan Krios | FEI Titan Krios |
| Voltage (keV) | 300 |  |  |  |  | 300 | 300 |
| Camera | Gatan K2-Summit |  |  |  |  | Gatan K2-Summit | Gatan K2-Summit |
| Electron exposure (e-/Å<sup>2</sup>) | 1.42 |  |  |  |  | 1.33 | 1.33 |
| Energy filter slit width (eV) | 20 (Gatan Quantum-LS (GIF)) |  |  |  |  | 20 (Gatan Quantum-LS (GIF)) | 20 (Gatan Quantum-LS (GIF)) |
| Pixel size (Å) | 0.82 |  |  |  |  | 0.64 | 0.64 |
| Defocus range (µm) | (-0.8) - (-2.8) |  |  |  |  | (-0.8) - (-2.8) | (-0.8) - (-2.8) |
| Magnification (nominal) | 60'975x (165kx) |  |  |  |  | 78'125x (215kx) | 78'125x (215kx) |
| Number of frames per movie | 35 |  |  |  |  | 45 | 45 |
| Number of good micrographs | 12'060 |  |  |  |  | 3'964 | 1'578 |
| Refinement procedure | As monomer |  |  | As dimer |  | As monomer | As dimer |
|  | Overall | Focused on C-terminal | Open conformation | Overall | Focused on dimerization interface |  |  |
| Initial particles (Before/After 2D cleaning) | 815'057/490'743 |  |  | 534'661/175'411 |  | 1'395'292/196'182 | 112'358/32'079 |
| Final particles | 184'206 | 184'206 (softmask) | 93'151 | 80'140 | 80'140 (softmask) | 119'115 | 32'079 |
| Map resolution (Å) | 3.40 | 3.43 | 4.72 | 5.27 | 3.93 | 4.60 | 7.46 |
| FSC threshold | 0.143 | 0.143 | 0.143 | 0.143 | 0.143 | 0.143 | 0.143 |
| Map resolution range (Å) | 20 - 2.60 | 20 - 2.90 | 20 - 4.10 | 20 - 4.30 | 20 - 3.40 | 20 - 4.0 | 20 - 6.50 |

**Suppl. table 2:** Summary of data collection and refinement statistic for the PMb21 structure obtained by X-ray crystallography. (*) numbers in brackets denote highest resolution shell.

|  |  |
| --- | --- |
|  | **PMb<sub>21</sub>** |
| **Data collection** |  |
| Number of crystals | 1 |
| Space group | I2 |
| Cell dimensions |  |
| a, b, c (Å) | 121.76 87.62 51.90 |
| $\alpha, \beta, \gamma$ (°) | 90.00 105.82 90.00 |
| No. of reflections | 248315 |
| No. of unique reflections | 34750 |
| Resolution (Å) | 44.25-2.00 (2.05-2.00)* |
| R<sub>merge</sub> | 0.057 (1.167) |
| R<sub>pim</sub> | 0.023 (0.469) |
| CC<sub>1/2</sub> | 0.999 (0.722) |
| I/σI | 15.1 (1.5) |
| Completeness (%) | 98.1 (96.7) |
| Redundancy | 7.1 (7.0) |
| **Refinement** |  |
| Resolution (Å) | 2.0 |
| No. of reflections | 34737 |
| R<sub>work</sub>/R<sub>free</sub> | 0.193/0.239 |
| No. atoms |  |
| Protein | 3766 |
| Ligand | 23 |
| Water | 153 |
| Other | 25 |
| B-factors |  |
| Protein | 62.6 |
| Ligand | 47.2 |
| Water | 54.3 |
| Other | 63.4 |
| R.m.s deviations |  |
| Bond length (Å) | 0.010 |
| Bond angle (°) | 1.021 |
| Ramachadran plot statistics (%)* |  |
| Favoured regions | 97.51 |
| Allowed regions | 2.29 |
| Disallowed regions | 0.21 |

## Methods

### Expression and purification of the *Neisseria gonorrhoeae* LptDE complex (NgLptDE)

*N.gonorrhoeae* wild type full length LptD (UniProt: Q5F651) and LptE (UniProt: Q5F9V6), with a hexhistidine tag on the c-terminus, were cloned into the expression vector pBAD22A (graciously provided by Professor Yihua Huang). NgLptDE complex was co transformed in SF100 *E.coli* cells (graciously provided by Professor Yihua Huang). After transformation, pre-culture was started, and cells were grown overnight at 37°C with 100μg/ml ampicilin and 50μg/ml kanamycin. Large LB broth culture was then inoculated at a starting OD_600nm_=0.05 and grown at 37°C with 100 μg/ml ampicillin. When OD_600nm_=0.8, temperature was switched to 20°C and induction was performed at OD_600nm_=1 with 0.4 % arabinose. After overnight induction at 20°C, cells were harvested and resuspended in lysis buffer consisting of 200 mM NaCl, 50 mM Tris-HCl pH 7.5, 10 mM MgCl_2_, 10 μg/ml DNAse I, 100 μg/ml AEBSF [4-(2-aminoethyl)benzenesulfonyl fluoride hydrochloride] and protease inhibitors cocktail (Complete, Roche). Cells were cracked by multiple passages through a microfluidizer system using a pressure of 18’000 psi, and the lysate was centrifuged at 7’500 x g for 10 minutes to remove the cell debris. The supernatant was collected, and inner membranes were solubilized by incubation with 2% Triton X-100 for 30 minutes at 4°C under gentle agitation. The outer membrane fraction was collected by centrifugation at 100’000 x g for 30 minutes at 4°C. The pellet containing the outer membrane fraction was resuspended in 200 mM NaCl, 50 mM Tris-HCl pH 7.5, 20 mM imidazole, 1 % LDAO [lauryldimethylamine oxide], supplemented with protease inhibitors cocktail (Complete, Roche) and incubated under gentle agitation for 12 hours at 4°C. Insoluble material was removed by centrifugation at 100’000 x g for 1 hour at 4°C. The supernatant was incubated in batch with ~ 5 ml NiNTA resin for 2 hours at 4°C under gentle agitation. The resin was subsequently washed by gravity flow with 10 column volumes (CV) of Wash buffer A (200 mM NaCl, 20 mM Tris-HCl pH 8.0, 20 mM imidazole, 1 % LDAO), 10 CV of Wash buffer B (150 mM NaCl, 20 mM Tris-HCl pH 8.0, 40 mM imidazole, 0.5 % LDAO), 10 CV of Wash buffer C (150 mM NaCl, 20 mM Tris-HCl pH 8.0, 40 mM imidazole, 0.2 % LDAO) and 10 CV of Wash buffer D (150 mM NaCl, 20 mM Tris-HCl pH 8.0, 40 mM imidazole, 0.1 % LMNG [lauryl maltose neopentyl glycol]). Elution was performed with 5 CV of Elution buffer (150 mM NaCl, 20 mM Tris-HCl pH 8.0, 300 mM imidazole, 0.01 % LMNG). Eluted material was desalted against 150 mM NaCl, 20 mM Tris-HCl pH 8.0, 0.005% LMNG using disposable PD-10 desalting columns (GE Healthcare) and was concentrated in 100 kDa centrifugal concentrators (Millipore). Concentrated sample was further purified by size exclusion chromatography on a Superdex 200 Increase column, equilibrated to 150 mM NaCl, 20 mM Tris-HCl pH8.0 and 0.005 % LMNG. The peak fractions corresponding to the monomeric or dimeric LptDE complex were kept separately, concentrated to ~ 1 mg/ml and use for sybody generation, as well as biophysical and structural studies.

### Protein used for sybodies generation and grating coupled Interferometry characterization

*N. gonorrhoeae* (Zopf) Trevisan (ATCC 700825) LptD (1-801)-3C-His10 and LptE (1-159)-AVI complex was produced using *E.coli* SF100 cells and purified as described above and an *in-vitro* biotinylation step was added after the size exclusion chromatography (SEC) step ^35^. Briefly, LptDE protein was concentrated to 10 μM concentration and mixed with 40 μg of BirA *E.coli* enzyme, 5 mM ATP, 10 mM MgAc and 15 μM biotin. The mixture was incubated 16 hours at 4°C and a second SEC step was performed to desalt the sample and remove the BirA enzyme and free Biotin in the following buffer: 20 mM Tris-HCl pH 8.0, 150 mM NaCl, 0.005 % LMNG. Fractions corresponding to the monomeric complex peak were pooled, concentrated and flash frozen in liquid nitrogen for subsequent use.

### Sybody generation

Sybodies were generated as described previously ^24,36^ with one notable difference. NgLptDE could not be produced in large amounts, which did not allow to use a large excess of non-biotinylated NgLptDE for off-rate selection. Therefore, we pooled all the sybodies present after the first round of phage display and used them as competitors during the second round of phage display to perform an off-rate selection. To this end, the pDX_init plasmid outputs (of the concave, loop and convex library) of the first round of phage display were purified by miniprep (Qiagen). FX cloning was performed to transfer the sybody pool from the pDX_init to the pSb_init expression plasmid using 2 μg of pDX_init pool and 1 μg of pSb_init. The cloning reaction was subsequently transformed into electrocompetent *E. coli* MC1061 cells (> 10 mio cfu). The sybody pools were expressed as described for single sybodies using the pSb_init construct ^36^. After expression of the pools in 600 ml cultures of TB medium, the sybodies were extracted from the cells by periplasmic extraction, purified by IMAC and dialyzed overnight against Tris Buffered Saline (TBS). Precipitation was removed by centrifugation at 20’000 x g for 15 min. The pools were used at a concentration of approximately 100 μM to perform an off-rate selection for 2 min in the second round of phage display.

### Fluorescent labelling of sybodies

To perform site specific labelling of the sybodies, a glycine located between SapI restriction site and myc tag on the pSb_init backbone was mutated to cysteine via Quick change mutagenesis, thereby adding the following amino acids to the C-terminus of the sybody: GRACEQKLISEEDLNSAVDHHHHHH. The sybodies were expressed and purified as previously described ^36^ except that 1 mM DTT was added to all buffers used for purification. Subsequently, DTT was removed and the sybody was re-buffered to degassed PBS using a PD10 desalting column and immediately mixing the sybody with Alexa Fluor 647 C_2_ maleimide (ThermoFisher Scientific) at a molar ratio of 1:3.6. The labelling reaction was carried out for 1 hour at 4°C. Excess label was removed by desalting the labelled sybody with a PD10 column.

### Cellular binding assay

For cellular binding assays, overnight cultures of *E. coli* SF100 cells with and without overexpression of NgLptDE were used. The number of cells were normalized by adjusting 1ml of culture to an OD_600_ of 3. The cells were harvested by centrifugation washed three times with 500 μl PBS containing 0.5 % BSA (PBS-BSA) and subsequently blocked for 20 min in the same buffer. After an additional wash with 500 μl PBS-BSA, the cells were incubated for 20 min in 100 μl PBS-BSA containing 1 μM of the Alexa Fluor 647-labelled sybodies. After three washes with 500 μl PBS, cells were resuspended in 100 μl PBS and transferred to a microtiter plate with non-transparent walls. Fluorescence was measured in a plate reader with excitation of 651 nm and emission of 671 nm.

### Antibiotic susceptibility assay

*N. gonorrhoeae* (Zopf) Trevisan (ATCC 700825) were streaked from a glycerol stock on blood agar and incubated for 24 hours at 37 °C with 5 % CO_2_ atmosphere. Colonies were scraped off the agar and resuspended in Fastidious broth at a density of McFarland 0.5 ^37^. The cells were further diluted 1:100 in Fastidious broth. In 96 well plates, dilution series of vancomycin with and without sybodies in Fastidious broth were prepared and mixed with the diluted culture. The plates were incubated without shaking at 37 °C with 5 % CO_2_ atmosphere for 24 hours. 100 μl of a 0.04 mg/ml resazurin stock solution in PBS was added to the cells and incubated for one hour at 37 °C with 5 % CO_2_ atmosphere. Fluorescence was measured at 571 nm excitation and 585 nm emission.

### Pro-Macrobody generation

Pro-Macrobodies (PMbs) were produced in *E.coli* as described earlier for the original macrobodies ^20^. Briefly, two PCR-amplified fragments, the specific sybody and the C-terminal MBP were cloned in parallel into the expression vector pBXNPH3M (Addgene #110099) ^38–40^ using FX cloning ^41^. N-terminally of the resulting PMb insert, the plasmid expresses a pelB leader sequence followed by a deca-His tag, an MBP and a 3C site. The insert is fused during cloning through an overlapping proline-encoding CCG codon introduced by reverse and forward primers at the 3’- and 5’-end of the sybodies and MBP respectively and released by digestion with the Type IIS restriction enzyme SapI (NEB). The second proline of the linker is encoded in the forward primer of the MBP (3’ of the overlapping CCG codon) and replaces the natural lysine. The resulting amino acid sequence of the linker is VTV***PP***<u>LVI</u> (VTV is the conserved C-terminus of sybodies, PP in italics/bold denotes the linker and underlined the truncated N-terminus of processed *E.coli* malE starting at Leu7). PMbs were expressed in terrific broth in MC1061 *E.coli* cells at 37°C by induction with 0.02% arabinose at an OD_600_=0.7. After 3.5 hours cells were harvested and resuspended in lysis buffer consisting of 150 mM NaCl, 50 mM Tris-HCl pH 8, 20 mM imidazole, 5 mM MgCl_2_, 10% glycerol, 10 μg/ml DNAse I and protease inhibitors (Complete, Roche). Cells were cracked and the lysate was centrifuged for 30 min at 147’000 x g in a Beckman 45Ti rotor. Subsequently, the supernatant was incubated in batch with ~ 3ml NiNTA resin/ 6L of culture. The resin was washed with 150 mM KCl, 40 mM imidazole pH7.6, 10% glycerol and eluted with 150 mM KCl, 300 mM imidazole pH 7.6, 10% glycerol. The N-terminal MBP with the deca-His tag was removed by cleavage with 3C protease overnight during dialysis against 150 mM KCl, 10 mM Hepes-NaOH, 20 mM imidazole, pH 7.6, 10% glycerol. After removal of the His-tagged MBP by Re-IMAC the unbound material was concentrated in 50 kDa centrifugal concentrators (Millipore) and subjected to SEC on a Superdex 200 Increase column (equilibrated 150 mM NaCl, 10 mM Hepes-Na, pH7.6). The eluted fractions with the PMbs were supplemented with 20% glycerol, concentrated to 3-8 mg/ml and aliquots were flash-frozen in liquid nitrogen for subsequent use. Macrobody versions of sybodies 21 and 51 with the original VK-linker were expressed and purified in the same way.

### SEC analysis/ purification of LptDE-PMb complexes

To identify ternary and quaternary complexes of NgLptDE with various PMbs against NgLptDE, monomeric NgLptDE was first separated from dimers by SEC on a Superdex 200 Increase 10/300 column in 150 mM NaCl, 20 mM Tris-Cl, pH8, 0.005% LMNG. A complex was formed by addition of 3 to 4-fold molar excess of PMb51 to NgLptDE sample. The uncomplexed and PMb-bound samples were analyzed on an Agilent 1260 Infinity II HPLC using a Superdex 200 Increase 5/150 column and Trp-fluorescence detection after 10-15 min incubation time on ice. Binding was indicated by a shift to earlier elution volumes and increase in the UV-absorption and Trp-fluorescence. The quaternary complex was formed by adding PMb21 in 3 to 4-fold molar excess to the preformed NgLptDE-PMb51 complex. The quaternary NgLptDE-PMb51-PMb21 complex showed the most distinct shift and fluorescence increase and was therefore chosen for scale up and cryo-EM analysis.

### Binding analytics by Grating Coupled Interferometry

Initial screen with ELISA positive sybodies was performed at 20°C on the Wave delta instrument from Creoptix in Tris 25mM pH 7.5, NaCl 300 mM, DDM 0.1% as running buffer. Biotinylated protein was immobilized on a 4PCP-S (streptavidin) chip, conditioned with 1M NaCl, 0.1 M sodium borate, at levels between 600 and 700 pg/mm2 to avoid any mass transport limitation. One injection of each sybody (200nM) was performed and binding responses were evaluated with the Wave control software. The ten best sybodies in term of slower K_off_ as well as quality of binding signals obtained were characterized further with dose-response analysis in order to determine accurate kinetic parameters. For each ten sybody, 8 concentrations were recorded (serial 2-fold dilution) in duplicate injections and equilibrium as well as kinetic data were analyzed and fitted with 1:1 model. Fits were of high quality (black curves) and recapitulate the experimental data (red curves). At higher concentration of sybodies, bulk (RI) effects could be observed, but this effect did not influence data analysis. PMb21 showed a slightly decreased affinity (3 to 4-fold) compared to the respective sybody 21. This loss can be explained by a faster K_off_ of the PMb21.

### Sample preparation and Cryo-EM data acquisition

Quantifoil (1/2) 200-mesh copper grids were glow-discharged for 20 seconds prior to sample freezing. 3μl of NgLptDE-PMb51-PMb21 complex at a concentration of 1mg/ml were placed on the grid, blotted for 3.0s and flash frozen in a mixture of liquid propane and liquid ethane cooled with liquid nitrogen using a Vitrobot Mark IV (FEI) operated at 4°C and under 100% humidity.

The EM data collection statistics in this study are reported in the **Suppl. table 1**. Data were recorded on a FEI Titan Krios transmission electron microscope, operated at 300 kV and equipped with a Quantum-LS energy filter (slit width 20 eV; Gatan Inc.) containing a K2 Summit direct electron detector. Data were automatically collected using the software SerialEM ^42^. Dose-fractionated exposures (movies) were recorded in electron-counting mode, applying 60 electrons per square Angstrom (e^−^/Å^2^) over 45 frames, or 50 e^−^/Å^2^ over 35 frames for, respectively, the NgLptDE (apo) or the NgLptDE-PMb51-PMb21 samples. A defocus range of −0.8 to −2.8 μm was used and the physical pixel size was 0.64 Å/pixel for the NgLptDE and 0.82 Å/pixel for the NgLptDE-PMb51-PMb21 datasets. Recorded data were online analyzed and pre-processed using FOCUS ^43^, which included gain-normalization, motion-correction, and calculation of dose-weighted averages with MotionCor2 ^44^, as well as estimation of micrograph defocus with CTFFIND4 ^45^.

### Image processing

The following processing workflows were used for the samples in the study. The aligned movies were imported into CryoSPARC V2 ^46^. A set of aligned averages with a calculated defocus range of −0.6 to – 3.0 μm was selected from which averages with poor CTF estimation statistics were discarded. Automated particle picking in CryoSPARC V2 resulted in 815,057 particle locations for the NgLptDE-PMb51-PMb21 sample. After several rounds of 2D classification, 490,743 particles were selected and subjected to 3D classification using the multi class *ab initio* refinement process (5 classes; 0.4 similarity) and heterogenous refinement. The best resolved class consisting of 184,206 particles was finally subjected to 3D non-uniform refinement. The overall resolution of the resulted map was estimated at 3.40 Å based on the Fourier shell correlation (FSC) at 0.143 cutoff ^47^. To visualize the LptD-NTD dimerization interface, another round of 3D heterogenous refinement was performed and a subset consisting of 80,140 particles was selected. Those particles coordinates were used re-extraction of particle images with an increased box size. Particles were re-centered by 2D classification in order to process the dimeric LptDE complex. *Ab initio* reconstruction and non-uniform refinement on this set of particles resulted in a _DIMER_NgLptDE-PMb51-PMb21 map with an overall resolution of 5.27 Å. As for the opened state, a multi class *ab initio* refinement process and heterogenous refinement was performed for the NgLptDE-PMb51-PMb21 sample. Of the five 3D classes, one class consisting of 93,151 particles was further refined by computationally removing the density corresponding to the detergent micelle with the particle subtraction tool within CryoSPARC V2, followed by 3D local refinement. The resulting map had an estimated overall resolution of 4.72 Å as judged by FSC at 0.143 cutoff. Analysis of the apo NgLptDE was performed similarly to the NgLptDE-PMb21-PMb51. Briefly, a set of aligned averages with a calculated defocus range of −0.6 to –3.0 μm was selected, from which averages with poor CTF estimation statistics were discarded. Automated particle picking in CryoSPARC V2 resulted in 1,395,392 particle locations for the NgLptDE. After several rounds of 2D classification, 196,182 particles were selected and subjected to 3D classification using the multi class *ab initio* refinement process (3 classes; 0.1 similarity) and heterogenous refinement for the 2 best classes. The best resolved class consisting of 119,115 particles was finally subjected to 3D non-uniform refinement. The overall resolution of the resulted map was estimated at 4.6 Å based on the Fourier shell correlation (FSC) at 0.143 cutoff.

### Model building and refinement

An initial LptDE model was generated using SWISS-MODEL ^48^, using as templates the KpLptD structure (PDB-ID 5IV9) and the EcLptE structure (PDB-ID 4RHB). The template for building the PMb_21_ coordinates into the EM map was based on the X-ray structure that was solved specifically for this study. The same structure was also used to model the maltose binding protein region of PMb_51_.

Rigid body fitting was initially done in Chimera ^49^ followed by manually rebuilding of the model in Coot ^50^. Remaining clashes between sidechains were detected using Schrodinger version 2019-4 (Maestro, Schrödinger, LLC, New York, NY, 2021), and remodelled using prime ^51^. Manual inspection of missing H-bonds in the model was used to refine sidechain positions. Finally, real-space refinement was performed in Phenix version 1.17-3644, applying Ramachandran plot restraints ^52^.

### Molecular dynamics simulations

Possible linkers connecting the VHH to MBP were assessed by all-atom MD simulations. After 300 ns of equilibration, conformational space was explored by conducting 500 ns MD trajectories initiated from a structure solved with a (Val122 and Lys123) VL-linker (PDB-ID 6HD8), as well as the same set of coordinates but modelled with an engineered (Pro122-Pro123) PP-linker. Both trajectories were later aligned on the VHH structure (residues 1-120) to show the larger conformational flexibility of the VK-linker. The macrobody interdomain (VHH to MBP) angle was also monitored during MD trajectories. This angle was defined as the angle between the geometrical centers of residue 1-120 (Nb), residue 122-123 (linker), and residue 124-486 (MBP). All MD simulations were conducted in explicit water, at room temperature (300K, or 26.85°C), and with Desmond standard parametrization ^53^ and the OPLS3e force field ^54^.

### Crystallization and structure determination of PMb_21_

Purified PMb_21_ in 150mM NaCl, 10 mM HEPES-Na, pH7.5, concentrated to 10 mg/ ml and supplemented with 2.5mM D-(+)-maltose was subjected to crystallization screens. The protein crystallized in 0.2 M MgCl_2_, 0.1M Hepes pH 7.5, 30% PEG400. For cryoprotection crystals were soaked in 0.2M MgCl_2_, 0.1M Hepes pH 7.5, 35% PEG400 and 2.5mM D-(+)-maltose. Crystals were flash frozen in liquid N_2_ and data was collected at the PXII (X10SA) beamline of the Swiss Light Source (SLS, Villigen). The dataset was integrated and scaled with XDS (built=20190806). The structure was solved by PHASER version 2.8.3 (CCP4 7.0.077) using MBP (PDB 1N3W) and a nanobody (PDB 5FWO) as molecular replacement search model. Structure was initially refined using REFMAC version 5.8.0257 (CCP4 7.0.077) and later with PHENIX version 1.16, together with iterative model building in COOT using 2FoFc and FoFc map.

### Figure preparation

Figures were prepared using the programs Chimera X (http://www.rbvi.ucsf.edu/chimerax/ ^55^), Chimera (http://www.cgl.ucsf.edu/chimera/ ^49^), PyMOL (http://www.pymol.org/) and Schrödinger (www.schrodinger.com).

